# Thiol reductive stress activates the hypoxia response pathway

**DOI:** 10.1101/2023.03.06.531274

**Authors:** Ravi, Ajay Kumar, Shalmoli Bhattacharyya, Jogender Singh

## Abstract

Owing to their capability to disrupt the oxidative protein folding environment in the endoplasmic reticulum (ER), thiol antioxidants such as dithiothreitol (DTT) are used as ER-specific stressors. We recently showed that thiol antioxidants modulate the methionine-homocysteine cycle by upregulating an S-adenosylmethionine-dependent methyltransferase, *rips-1*, in *Caenorhabditis elegans*. However, the changes in cellular physiology induced by thiol stress that modulate the methionine-homocysteine cycle remain uncharacterized. Here, using forward genetic screens in *C. elegans*, we discover that thiol stress enhances *rips-1* expression via the hypoxia response pathway. We demonstrate that thiol stress activates the hypoxia response pathway. The activation of the hypoxia response pathway by thiol stress is conserved in human cells. The hypoxia response pathway enhances thiol toxicity via *rips-1* expression and confers protection against thiol toxicity via *rips-1*-independent mechanisms. Finally, we show that DTT might activate the hypoxia response pathway by producing hydrogen sulfide. Our studies reveal an intriguing interaction between thiol-mediated reductive stress and the hypoxia response pathway and challenge the current model that thiol antioxidant DTT disrupts only the ER milieu in the cell.

## Introduction

Maintenance of cellular homeostasis requires a balanced redox environment in the cell. Redox reactions result in the production of reactive oxygen species (ROS). High levels of ROS are associated with several pathological conditions (Sies & Jones, 2020). However, physiological ROS plays essential roles in cellular survival, differentiation, proliferation, repair, and aging (Trachootham *et al*, 2008; Sies & Jones, 2020). An increased amount of antioxidants results in the depletion of ROS, leading to reductive stress that is linked with several pathological conditions (Pérez-Torres *et al*, 2017; Rajasekaran, 2020). Dietary supplementation of high amounts of antioxidants correlates with accelerated cancer progression and a higher incidence of cancer-related mortality (Villanueva & Kross, 2012; Sayin *et al*, 2014). A high amount of dietary antioxidants accelerates aging in the nematode *Caenorhabditis elegans* (Gusarov *et al*, 2021). Despite the association of high amounts of antioxidants with pathological conditions, the full spectrum of physiological effects of reductive stress remains to be elucidated.

Thiols, which contain a sulfhydryl group, are a major defense mechanism in the cell against oxidative stress and undergo oxidation to form disulfides under oxidative conditions (Ulrich & Jakob, 2019). Alterations in thiol-disulfide homeostasis play a role in several pathological conditions, including cardiovascular disease (Kundi *et al*, 2015), chronic renal disease (Rodrigues *et al*, 2013), and diabetes mellitus (Ates *et al*, 2016). However, the physiological effects of thiol reductive stress remain obscurely characterized. Because disulfide bonds are formed in the oxidative environment of the endoplasmic reticulum (ER), ER is expected to be the primary target of reductive stress (Cuozzo & Kaiser, 1999; Merksamer *et al*, 2008). Indeed, thiols such as dithiothreitol (DTT) disrupt disulfide bonds in the ER, leading to protein misfolding (Braakman *et al*, 1992). The ensuing protein misfolding results in ER stress; therefore, thiol reducing agents are used as ER-specific stressors (Oslowski & Urano, 2011). In addition to ER stress, thiol antioxidants can also result in oxidative stress by activating futile oxidative cycles in the ER (Maity *et al*, 2016). Recent studies in *C. elegans* showed that, besides causing ER stress, thiol antioxidants modulate the methionine-homocysteine cycle (Gokul & Singh, 2022; Winter *et al*, 2022). The modulation of the methionine-homocysteine cycle is a consequence of the upregulation of an S-adenosylmethionine (SAM)-dependent methyltransferase, *rips-1*, by thiol antioxidants. Thus, reductive stress caused by thiols could affect cellular physiology beyond the ER. However, such physiological effects of thiol antioxidants remain to be fully characterized.

In this study, using *C. elegans* and human cell lines, we characterized the physiological effects of thiol reductive stress. Because thiol antioxidants result in the increased expression of *rips-1* (Gokul & Singh, 2022; Winter *et al*, 2022), we reasoned that understanding the regulation of *rips-1* expression might provide insights into the physiological effects of thiol reductive stress. To this end, we carried out forward genetic screens to understand the regulation of *rips-1* expression. The expression levels of *rips-1* correlated with the activity of the hypoxia response pathway. We observed that exposure to DTT resulted in the activation of a functional hypoxia response pathway that protected *C. elegans* against cyanide-mediated fast killing on *Pseudomonas aeruginosa* PAO1. The hypoxia response pathway has *rips-1*-dependent and independent opposing roles in regulating thiol-mediated toxicity. The activation of the hypoxia response pathway by DTT exposure is conserved in human cell lines. We showed that DTT produces hydrogen sulfide, which might be responsible for activating the hypoxia response pathway. These studies reveal an intriguing relationship between thiol reductive stress and the hypoxia response pathway.

## Results

### The hypoxia response pathway modulates *rips-1* expression levels

The thiol antioxidants, DTT and β-mercaptoethanol, cause toxicity in *C. elegans* by upregulating a SAM-dependent methyltransferase, *rips-1*, that results in the modulation of the methionine-homocysteine cycle (Gokul & Singh, 2022; Winter *et al*, 2022). Previous studies have shown that *rips-1* expression might be altered by hydrogen sulfide, mitochondrial dysfunction, and the hypoxia-inducible factor (Nargund *et al*, 2012; Pender & Horvitz, 2018; Miller *et al*, 2011; Vora *et al*, 2022; Winter *et al*, 2022). However, the full spectrum of physiological changes induced by thiol reductive stress that results in *rips-1* upregulation remains to be discovered. Because thiol antioxidants activate the unfolded protein response (UPR) in the ER by enhancing protein misfolding, we asked whether the ER UPR pathways were involved in the upregulation of *rips-1* upon DTT exposure. To this end, we crossed a *rips-1p::GFP* reporter strain with mutants of different ER UPR pathways, including *ire-1(v33)*, *xbp-1(tm2482)*, *atf-6(ok551)*, and *pek-1(ok275)*. The green fluorescent protein (GFP) levels were lower in the *ire-1*/*xbp-1* and *atf-6* pathway mutants compared to the wild-type animals in the absence of DTT (Fig S1A and B). This could be because of compensatory protein translation inhibition by *pek-1* pathway due to increased ER stress in the *ire-1*/*xbp-1* and *atf-6* pathway mutants (Harding *et al*, 2000). It is also possible that the *ire-1*/*xbp-1* and *atf-6* pathways control the basal expression of *rips-1*. Nevertheless, exposure to DTT resulted in the upregulation of GFP levels in all the ER UPR mutants (Fig S1A and B), indicating that the ER UPR pathways are not involved in the DTT-mediated *rips-1* upregulation. Recently, we showed that DTT also results in the activation of the mitochondrial UPR (Gokul & Singh, 2022). Therefore, we asked whether the mitochondrial UPR is required for DTT-mediated upregulation of *rips-1*. DTT exposure resulted in the upregulation of *rips-1p::GFP* in *atfs-1(gk3094)* animals (Fig S1A and B), ruling out a role of mitochondrial UPR in the upregulation of *rips-1*.

To characterize the physiological changes that trigger the upregulation of *rips-1*, we designed a forward genetic screen to isolate mutants that had constitutive activation of *rips-1* (Fig S1C). We obtained three independent mutants with enhanced GFP expression compared to the parental strain JSJ13 (*rips-1p::GFP*) (Fig 1A and B). To identify the causative mutations, we performed whole-genome sequencing of the mutants after backcrossing them six times with the parental strain. The sequenced genomes of mutants were aligned with the reference genome of *C. elegans* to identify variants. Evaluation of the protein-coding genes having mutations in the isolated mutants revealed that all of the mutants had mutations in different negative regulators of the hypoxia response pathway (Fig 1C and Table S1). The JSJ14 mutant animals had two missense mutations in the gene *rhy-1*, the JSJ15 mutant animals had a splice site acceptor mutation in the gene *egl-9*, and the JSJ16 mutant animals had a premature nonsense mutation in the gene *vhl-1* (Fig 1C and Table S1). Because *egl-9(jsn15)* and *vhl-1(jsn16)* are likely loss-of-function alleles, these data suggested that inhibition of the negative regulators of the hypoxia response pathway may result in the enhanced expression of *rips-1*. Indeed, the knockdown of *egl-9*, *rhy-1*, and *vhl-1* by RNA interference (RNAi) resulted in the upregulation of *rips-1p::GFP* (Fig 1D and E).

**Figure 1.**
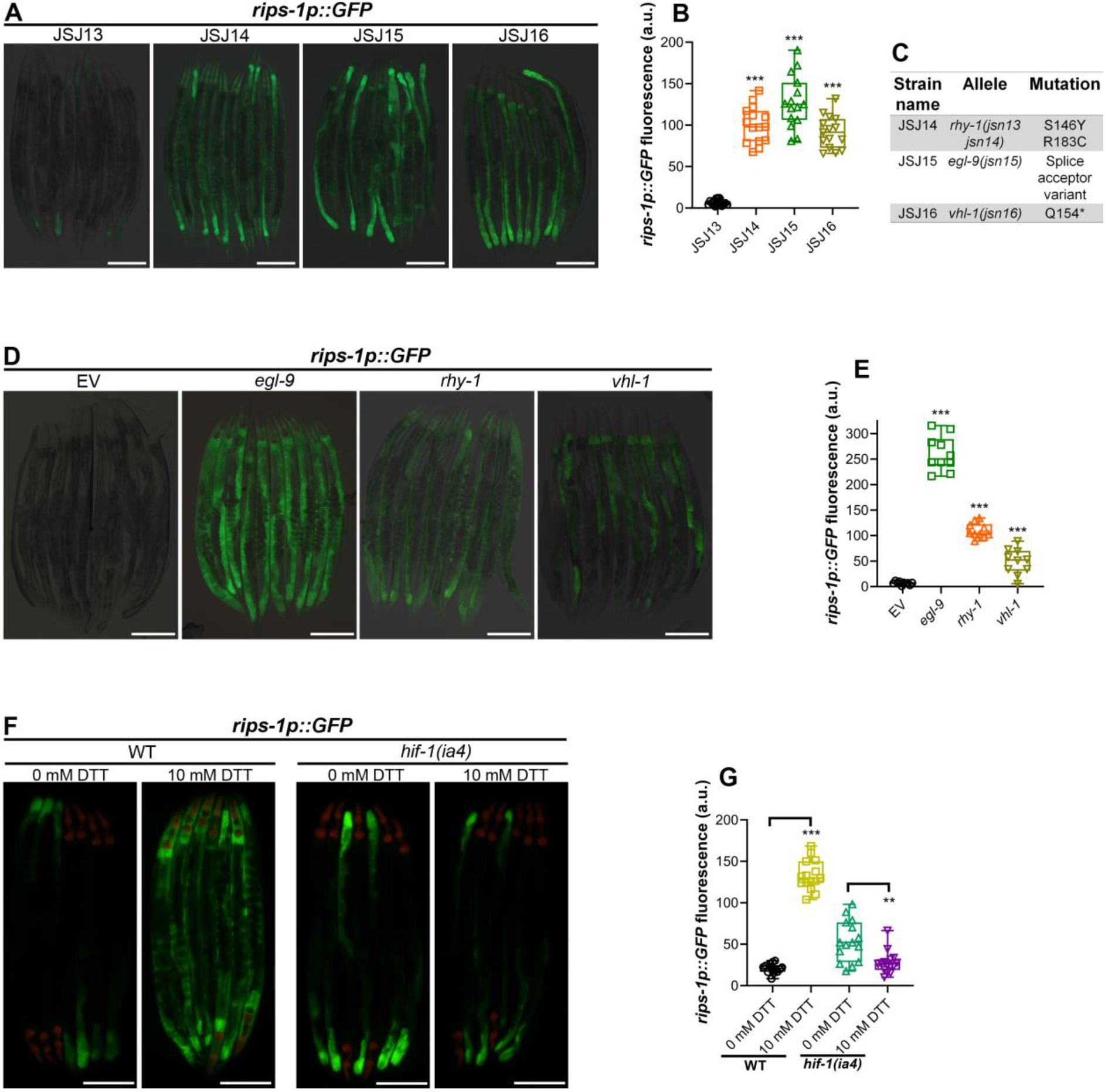
The hypoxia response pathway modulates *rips-1* expression levels. (A) Representative fluorescence images of the parental strain JSJ13 (*rips-1p::GFP*) and the mutants isolated from the forward genetic screen. The isolated mutants were designated JSJ14 to JSJ16. Scale bar = 200 µm. (B) Quantification of GFP levels of *rips-1p::GFP* in JSJ13-JSJ16 animals. ***p < 0.001 via the *t*-test (*n* = 15 worms each). (C) Table summarizing the alleles identified by whole-genome sequencing in JSJ14-JSJ16 strains. (D) Representative fluorescence images of *rips-1p::GFP* animals following RNAi against *egl-9*, *rhy-1*, and *vhl-1*. Animals grown on empty RNAi vector (EV) were used as the control. Scale bar = 200 µm. (E) Quantification of GFP levels of *rips-1p::GFP* animals following RNAi against *egl-9*, *rhy-1*, and *vhl-1*, along with EV control. ***p < 0.001 via the *t*-test (*n* = 10 worms each). (F) Representative fluorescence images of *rips-1p::GFP* and *hif-1(ia4);rips-1p::GFP* animals grown on 0 mM DTT until the young adult stage, followed by incubation on 0 mM or 10 mM DTT for 10 hours. The red fluorescence in the pharynx region is from the *myo-2p::mCherry* coinjection marker. Scale bar = 200 µm. (G) Quantification of GFP levels of *rips-1p::GFP* and *hif-1(ia4);rips-1p::GFP* animals grown on 0 mM DTT until the young adult stage, followed by incubation on 0 mM or 10 mM DTT for 10 hours. ***p < 0.001 and **p < 0.01 via the *t*-test (*n* = 14-15 worms each).

The genes *egl-9*, *rhy-1*, and *vhl-1* are involved in the degradation of the transcription factor hypoxia-inducible factor 1 (HIF-1) (Epstein *et al*, 2001; Shen *et al*, 2006). Inhibition of these genes results in the accumulation of HIF-1 and activates the expression of hypoxia response genes (Shen *et al*, 2006). We asked whether the upregulation of *rips-1* also required HIF-1 by using *hif-1(ia4)* animals. The *hif-1(ia4)* allele is a predicted null allele with deletion of exons 2–4 and lacks the induction of hypoxia response genes (Shen *et al*, 2005). The basal expression of *rips-1* was independent of HIF-1 as *hif-1(ia4);rips-1p::GFP* animals had GFP in the posterior region of the intestine (Fig 1F and G). Indeed, *hif-1(ia4);rips-1p::GFP* animals showed slightly increased levels of GFP in the posterior region of the intestine compared to *rips-1p::GFP* animals suggesting a compensatory induction of basal levels of *rips-1* in the absence of HIF-1. Next, we exposed the *hif-1(ia4);rips-1p::GFP* animals to DTT and monitored changes in GFP levels. While DTT exposure resulted in high GFP levels in *rips-1p::GFP* animals, GFP levels did not increase in *hif-1(ia4);rips-1p::GFP* animals (Fig 1F and G). Taken together, these data indicated that the hypoxia response pathway regulates the expression of *rips-1*.

### CYSL-1 is required for DTT-mediated *rips-1* upregulation

To better understand the interplay between *rips-1* expression and the hypoxia response pathway, we carried out another forward genetic screen to isolate mutants that failed to upregulate *rips-1* upon exposure to DTT—a phenotype similar to *hif-1* loss-of-function mutants (Fig S2A). We isolated two mutants that failed to upregulate *rips-1* upon exposure to DTT (Fig 2A-D). To identify the causative mutations, the variants in the mutant animals were identified by whole-genome sequencing. Evaluation of the protein-coding genes having mutations in the linked regions revealed that both mutants had a missense mutation in the gene *cysl-1* (Fig 2E and Table S1). Previous studies have identified *cysl-1* as a positive regulator of the hypoxia response pathway that inhibits *egl-9* activities (Ma *et al*, 2012; Wang *et al*, 2022).

**Figure 2.**
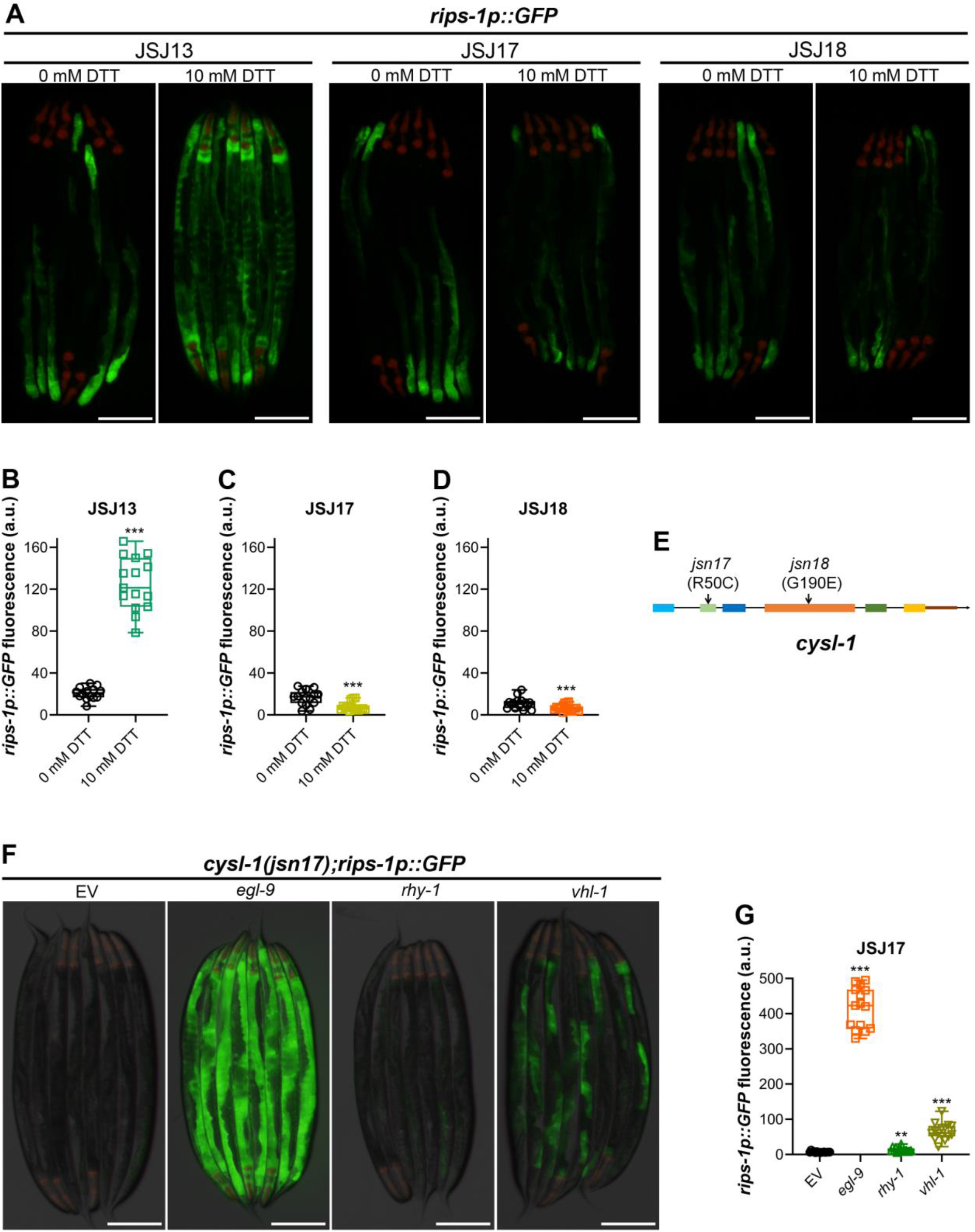
CYSL-1 is required for DTT-mediated *rips-1* upregulation. (A) Representative fluorescence images of the parental strain JSJ13 (*rips-1p::GFP*) and the isolated mutants, JSJ17 and JSJ18, grown on 0 mM DTT until the young adult stage, followed by incubation on 0 mM or 10 mM DTT for 10 hours. The red fluorescence in the pharynx region is from the *myo-2p::mCherry* coinjection marker. Scale bar = 200 µm. (B-D) Quantification of GFP levels of *rips-1p::GFP* in JSJ13 (B), JSJ17 (C), and JSJ18 (D) animals grown on 0 mM DTT until the young adult stage, followed by incubation on 0 mM or 10 mM DTT for 10 hours. 0 mM DTT data in Fig 2B is the same as 0 mM WT data in Fig 1G. ***p < 0.001 via the *t*-test (*n* = 14-15 worms each). (E) Mapping of the *cysl-1* alleles identified in the forward genetic screen. (F) Representative fluorescence images of JSJ17 (*cysl-1(jsn17);rips-1p::GFP*) animals following RNAi against *egl-9*, *rhy-1*, and *vhl-1*, along with EV control. Scale bar = 200 µm. (G) Quantification of GFP levels of JSJ17 (*cysl-1(jsn17);rips-1p::GFP*) animals following RNAi against *egl-9*, *rhy-1*, and *vhl-1*, along with EV control. ***p < 0.001 and **p < 0.01 via the *t*-test (*n* = 15 worms each).

Because *cysl-1* may act upstream or downstream of *hif-1* depending on the context (Budde & Roth, 2011; Ma *et al*, 2012), we carried out an epistasis analysis of *cysl-1* with the other components of the hypoxia response pathway in the context of *rips-1* expression. To this end, we knocked down *egl-9*, *rhy-1*, and *vhl-1* genes in *cysl-1(jsn17);rips-1p::GFP* and *cysl-1(jsn18);rips-1p::GFP* animals and monitored the changes in GFP expression. Knockdown of *egl-9* and *vhl-1* resulted in the upregulation of *rips-1* in both the alleles of *cysl-1* (Figs 2F and G, and S2B and C), indicating that *egl-9* and *vhl-1* do not require *cysl-1* to regulate the expression of *rips-*1 and work downstream of *cysl-1* to regulate *hif-1*. On the other hand, the knockdown of *rhy-1* did not lead to the upregulation of *rips-1* in *cysl-1* mutants (Figs 2F and G, and S2B and C), indicating *rhy-1* functions upstream of *cysl-1*. Therefore, in line with previous observations (Ma *et al*, 2012), these results indicated that *cysl-1* acts downstream of *rhy-1* and upstream of *egl-9, vhl-1*, and *hif-1* in *rips-1* expression.

### Thiol reductive stress activates the hypoxia response pathway

The involvement of the hypoxia response pathway in DTT-mediated upregulation of *rips-1* suggested that thiol reductive stress may activate the hypoxia response pathway. To test this possibility, we analyzed the mRNA levels of several genes that are induced by hypoxia, including *cysl-2*, *rhy-1*, *nhr-57*, *fmo-2*, *sqrd-1*, *gst-19*, and *oac-54* (Shen *et al*, 2005). All of these genes had significantly increased levels in DTT-exposed animals compared with the control animals (Fig 3A). The DTT-mediated induction of most of the hypoxia-response genes was dependent on *hif-1* (Fig 3B). The only exceptions were *nhr-57* and *fmo-2*. The upregulation of *nhr-57* was partially dependent on *hif-1*, while the upregulation of *fmo-2* was independent of *hif-1* (Fig 3A and B). The expression of *fmo-2* under hypoxic conditions is known to be regulated by the transcription factor NHR-49 (Doering *et al*, 2022). Further, previous studies have also reported that the expression of *nhr-57* is either partially or fully independent of HIF-1 (Shen *et al*, 2005; Doering *et al*, 2022). Taken together, these studies showed that DTT exposure results in the activation of a primarily HIF-1-mediated hypoxia response.

**Figure 3.**
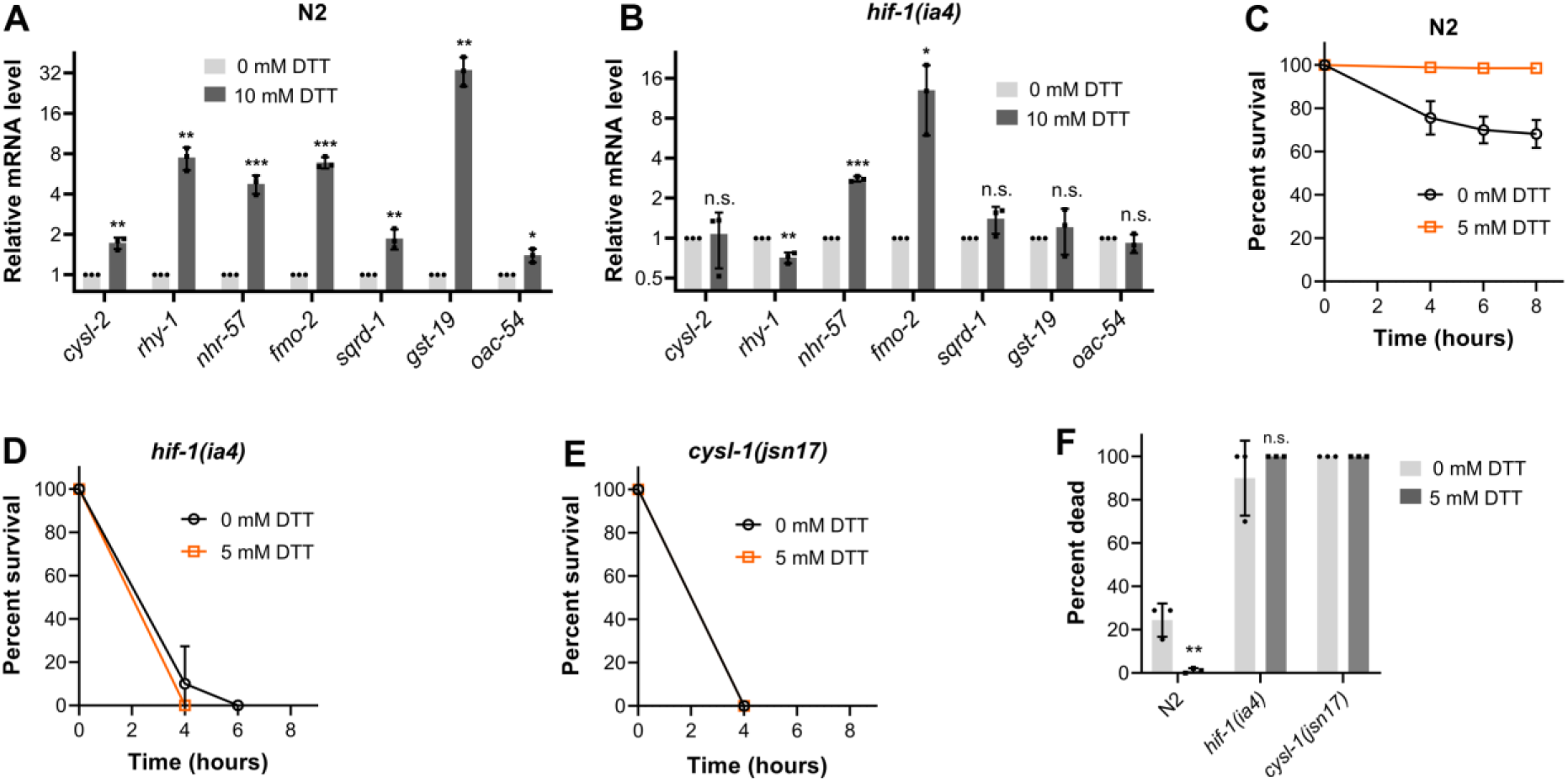
Thiol reductive stress activates the hypoxia response pathway. (A-B) Gene expression analysis of N2 (A) and *hif-1(ia4)* (B) animals grown on 0 mM DTT until the young adult stage, followed by incubation on 0 mM or 10 mM DTT for 10 hours. ***p < 0.001, **p < 0.01, and *p < 0.05 via the *t*-test. n.s., nonsignificant (*n* = 3 biological replicates). (C-E) Survival plots of N2 (C), *hif-1(ia4)* (D), and *cysl-1(jsn17)* (E) animals on *P. aeruginosa* PAO1 under fast-killing assay conditions. Before transferring to *P. aeruginosa* PAO1 lawns at the L4 stage, the animals were grown on *Comamonas aquatica* DA1877 diet containing 0 mM or 5 mM DTT. The plots of *cysl-1(jsn17)* animals are identical, as all the animals (from 0 mM and 5 mM DTT) are dead by the first time point (4 hours). Data represent mean and standard deviation from three independent experiments. (F) Percent dead animals on *P. aeruginosa* PAO1 under fast-killing assay conditions after 4 hours of exposure. Before transferring to *P. aeruginosa* PAO1 lawns at the L4 stage, the animals were grown on *Comamonas aquatica* DA1877 diet containing 0 mM or 5 mM DTT. Data represent mean and standard deviation from three independent experiments. **p < 0.01 via the *t*-test. n.s., nonsignificant.

Next, we asked whether DTT activated a functional hypoxia response pathway. Enhanced activity of the hypoxia response pathway has been shown to impart resistance to hydrogen cyanide-mediated fast paralytic killing of *C. elegans* by *P. aeruginosa* PAO1 (Gallagher & Manoil, 2001). We observed that pre-treatment with DTT enhanced the resistance of wild-type N2 animals to fast paralytic killing by *P. aeruginosa* PAO1 (Fig 3C). The hypoxia response pathway mutants exhibited enhanced susceptibility to *P. aeruginosa* PAO1 (Fig 3D and E), confirming the requirement of the hypoxia response pathway to protect against paralytic killing by *P. aeruginosa* PAO1. Importantly, pre-treatment with DTT did not enhance the survival of the hypoxia response pathway mutants (Fig 3D-F). These results indicated that DTT activates a functional hypoxia response pathway that protects *C. elegans* against hydrogen cyanide-mediated fast paralytic killing by *P. aeruginosa* PAO1.

### Hypoxia response pathway has opposing roles in regulating thiol-mediated toxicity

Next, we asked whether activation of the hypoxia response pathway had any role in regulating thiol-mediated toxicity. Thiol antioxidants-mediated upregulation of *rips-1* modulates the methionine-homocysteine cycle and results in developmental toxicity (Gokul & Singh, 2022; Winter *et al*, 2022). Therefore, it appears that the activation of the hypoxia response pathway by thiol antioxidants, which results in the upregulation of *rips-1*, might be harmful to the host. To understand the role of the hypoxia response pathway in toxicity by thiol reductive stress, we studied the response of *hif-1(ia4)*, *cysl-1(jsn17)*, and *cysl-1(jsn18)* animals to varying concentrations of DTT. Compared to wild-type N2 animals, *hif-1* and *cysl-1* loss-of-function mutants had improved development on 5 and 7.5 mM DTT (Figs 4A-D and S3A-C). These results indicated that the hypoxia response pathway exacerbated thiol toxicity, likely by increasing the expression of *rips-1*.

**Figure 4.**
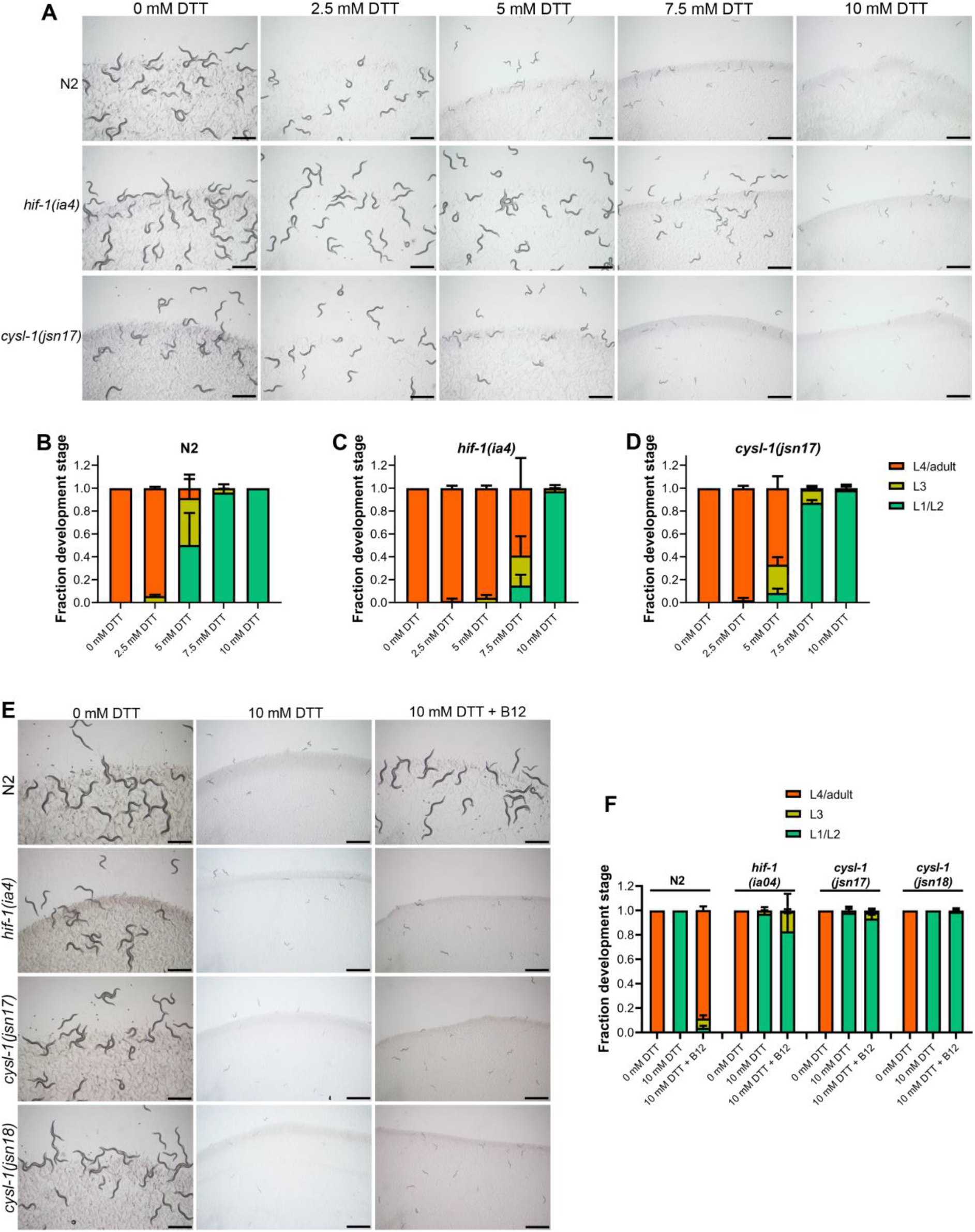
Hypoxia response pathway has opposing roles in regulating thiol-mediated toxicity. (A) Representative images of N2, *hif-1(ia4)*, and *cysl-1(jsn17)* animals on various concentrations of DTT on *E. coli* OP50 diet after 72 hours of hatching at 20°C. Scale bar = 1 mm. (B-D) Quantification of different developmental stages of N2 (B), *hif-1(ia4)* (C), and *cysl-1(jsn17)* (D) animals on various concentrations of DTT on *E. coli* OP50 diet after 72 hours of hatching at 20°C. (*n* = 3 biological replicates; animals per condition per replicate >100). (E) Representative images of N2, *hif-1(ia4)*, *cysl-1(jsn17)*, and *cysl-1(jsn18)* animals after 72 hours of hatching at 20°C on *E. coli* OP50 diet containing 0 mM DTT, 10 mM DTT, and 10 mM DTT supplemented with 50 nM vitamin B12. Scale bar = 1 mm. (F) Quantification of different developmental stages of N2, *hif-1(ia4)*, *cysl-1(jsn17)*, and *cysl-1(jsn18)* animals after 72 hours of hatching at 20°C on *E. coli* OP50 diet containing 0 mM DTT, 10 mM DTT, and 10 mM DTT supplemented with 50 nM vitamin B12. The 10 mM DTT is the same as in (B-D) (*n* = 3 biological replicates; animals per condition per replicate >100).

Interestingly, at 10 mM DTT, the hypoxia response pathway mutants failed to develop, similar to the wild-type animals (Figs 4A-D and S3A-C). Because vitamin B12 restores the DTT-perturbed methionine-homocysteine cycle, vitamin B12 alleviates DTT toxicity in wild-type animals (Gokul & Singh, 2022; Winter *et al*, 2022). Despite lacking induction of *rips-1*, the hypoxia response pathway mutants failed to develop on 10 mM DTT (Fig 4A-4D). Therefore, we asked whether supplementation of vitamin B12 would alleviate DTT toxicity in the hypoxia response pathway mutants. While the supplementation of 50 nM vitamin B12 alleviated DTT toxicity in wild-type N2 animals, it failed to do so in the hypoxia response pathway mutants (Fig 4E and F). Similarly, the wild-type animals developed on 10 mM DTT in the presence of the bacterial diet *Comamonas aquatica* DA1877, which has higher vitamin B12 compared to the *E. coli* OP50 diet (Gokul & Singh, 2022; Watson *et al*, 2014), while the hypoxia response pathway mutants failed to develop under these conditions (Fig S4A and B). Vitamin B12 did not affect the expression of *rips-1* either in the absence or presence of DTT (Fig S4C and D), indicating that vitamin B12 does not regulate thiol toxicity by modulating the hypoxia response pathway. Together, these results revealed a complicated picture of the role of the hypoxia response pathway in regulating thiol-mediated toxicity. By enhancing the expression of *rips-1* under thiol reductive stress, the hypoxia response pathway predisposes *C. elegans* to thiol-mediated toxicity. However, at higher concentrations of DTT, it emerges that the hypoxia response pathway protects against thiol reductive stress independent of *rips-1* expression as *hif-1* and *cysl-1* mutants fail to develop on 10 mM DTT even upon supplementation of vitamin B12.

### Activation of the hypoxia response pathway by thiol reductive stress is conserved in human cell lines

Next, we asked whether the activation of the hypoxia response pathway by thiol reductive stress is a conserved response. To this end, we used three human cell lines from different tissue origins, including HeLa (cervical cancer), LN229 (glioblastoma), and Cal27 (oral squamous cell carcinoma). First, we tested whether the survival of these cell lines was affected by exposure to 10 mM DTT. Exposure of these cell lines to 10 mM DTT for 24 hours did not affect their viability (Fig S5). We then evaluated the mRNA levels of five hypoxia-response genes, including hypoxia-inducible factor-1*α* (*HIF-1α), HIF-1β,* 3-phosphoinositide-dependent protein kinase-1 (*PDK1),* glucose transporter 1 *(GLUT1),* and BCL2/adenovirus E1B 19 kDa protein-interacting protein 3 (*BNIP3)* upon exposure of the cells to 10 mM DTT for 24 hours. All of these genes are known to be upregulated under hypoxic conditions (Choudhry & Harris, 2018; Guo *et al*, 2001; BelAiba *et al*, 2007). DTT treatment also resulted in the upregulation of these genes in HeLa, LN229, and Cal27 cell lines (Fig 5A-5C). These results showed that thiol reductive stress activates the hypoxia response pathway in human cell lines.

**Figure 5.**
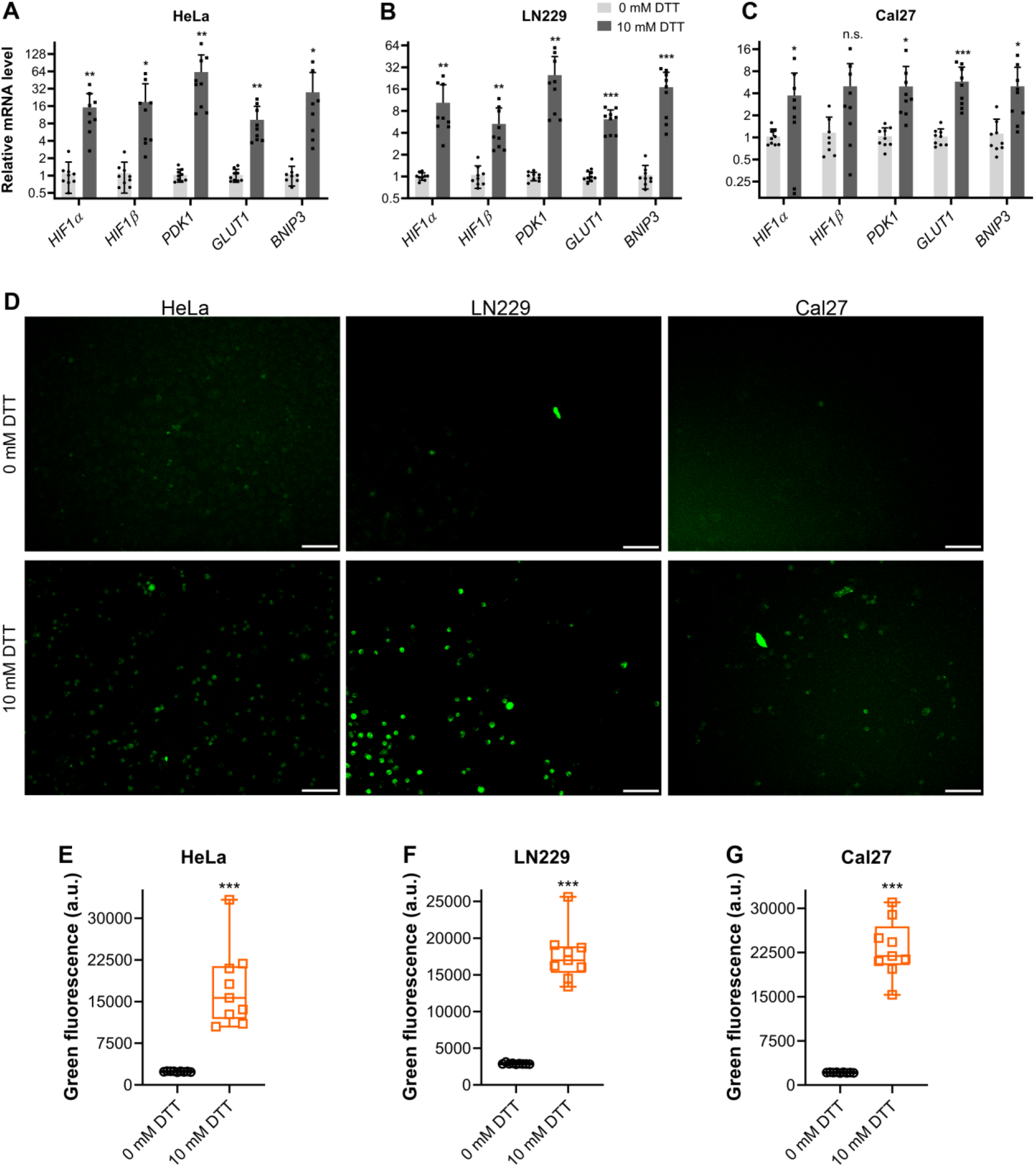
Thiol reductive stress activates the hypoxia response pathway in human cell lines. (A-C) Gene expression analysis of HeLa (A), LN229 (B), and Cal27 (C) cell lines with and without exposure to 10 mM DTT for 24 hours. ***p < 0.001, **p < 0.01, and *p < 0.05 via the *t*-test. n.s., nonsignificant (*n* = 3 biological replicates with 2-3 technical repeats each). (D) Representative fluorescence images of HeLa, LN229, and Cal27 cell lines upon treatment with Image-iT™ green hypoxia reagent. The cells were pretreated with 0 mM or 10 mM DTT for 24 hours before treatment with Image-iT™ green hypoxia reagent. Scale bar = 100 µm. (E-G) Quantification of fluorescence levels of Image-iT™ green hypoxia reagent-treated HeLa (E), LN229 (F), and Cal27 (G) cell lines. The cells were pretreated with 0 mM or 10 mM DTT for 24 hours before treatment with Image-iT™ green hypoxia reagent. ***p < 0.001 via the *t*-test (*n* = 9 wells each).

Next, we tested whether DTT resulted in a hypoxia-like cellular milieu. To this end, we stained human cell lines with Image-iT green hypoxia reagent after treatment with 10 mM DTT. Image-iT green hypoxia reagent is a fluorogenic compound that is live cell permeable and senses hypoxic environments. While it is unclear whether the Image-iT green hypoxia reagent senses oxygen levels directly or reports other hypoxia-related physiological changes, it fluoresces only under hypoxic conditions. DTT treatment resulted in a drastic increase in fluorescence in HeLa, LN229, and Cal27 cell lines (Fig 5D-G), indicating a hypoxia-like cellular milieu upon DTT exposure. To study whether the increase in the fluorescence was because of a hypoxic milieu or a change in the redox environment because of DTT, we tested the fluorescence changes in cells upon exposure to 10 mM *N*-acetylcysteine (NAC). Similar to DTT, NAC results in a reduced cellular environment, but it does not lead to the upregulation of *rips-1* (Gokul & Singh, 2022). The Image-iT hypoxia reagent fluorescence remained unaltered in cells exposed to 10 mM NAC for 24 hours (Fig S6), indicating that the increased fluorescence upon DTT exposure was unlikely because of a change in the redox environment.

### DTT leads to hydrogen sulfide production

Finally, we asked how thiol reductive stress led to the activation of the hypoxia response pathway. Thiol antioxidants result in the breakage of disulfide bonds in the ER (Maity *et al*, 2016). This is followed by the activation of futile oxidative cycles in the ER, enhancing disulfide bond turnover (Maity *et al*, 2016). Disulfide bond formation consumes oxygen and produces hydrogen peroxide (Zito, 2015). Therefore, thiol reductive stress generates hydrogen peroxide and causes oxidative stress (Maity *et al*, 2016). We hypothesized that thiol reductive stress might activate the hypoxia response pathway by creating oxidative stress or directly enhancing oxygen consumption. To test these possibilities, we probed the role of oxidative stress in the upregulation of *rips-1*. While oxidative stress by tert-butyl hydroperoxide, ferric chloride, and paraquat resulted in the induction of oxidative stress reporter *gst-4p::GFP*, it did not affect the expression of *rips-1p::GFP* (Fig S7A-D). These results suggested that oxidative stress is not responsible for the DTT-triggered hypoxia response pathway activation.

We then tested whether thiol reductive stress resulted in enhanced oxygen consumption. To this end, we measured oxygen consumption rates (OCR) in human cell lines upon exposure to 10 mM DTT. The addition of DTT resulted in a rapid increase in OCR not only in the cell lines but also in a blank control of medium without cells (Fig S7E-H). This indicated that the addition of DTT might increase OCR because of the oxidation of DTT, which results in oxygen consumption. However, the cells did not exhibit a further increase in OCR, indicating that DTT did not increase oxygen consumption in human cells. These results suggested that DTT does not activate the hypoxia response pathway by enhancing oxygen consumption.

Hydrogen sulfide (H_2_S) is known to activate the hypoxia response pathway via CYSL-1 (Ma *et al*, 2012). Because DTT also required *cysl-1* for the activation of the hypoxia response pathway (Fig 2), we asked whether DTT resulted in H_2_S production. To this end, we incubated worm lysate with DTT and measured H_2_S production. While the worm lysate in the absence of DTT did not lead to H_2_S production, H_2_S formation was observed in the presence of DTT (Fig 6A). The worm lysate was required for H_2_S production by DTT, as the addition of DTT to the buffer control did not result in the production of H_2_S (Fig 6A). Therefore, it appeared that DTT might result in the production of H_2_S in *C. elegans*.

**Figure 6.**
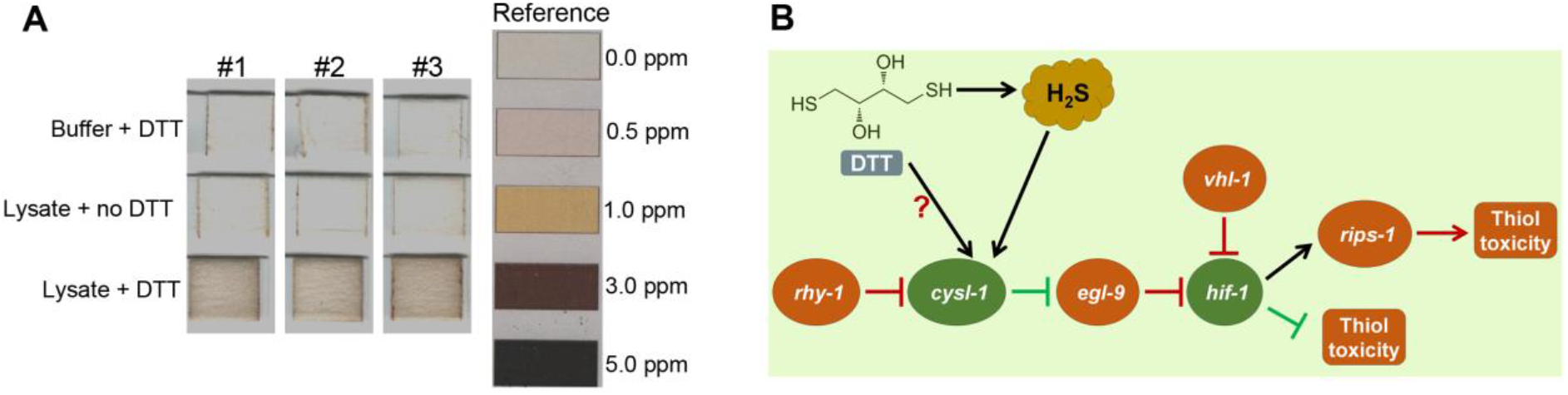
The mechanism of activation of the hypoxia response pathway by DTT. (A) Images of lead acetate paper showing H_2_S production in worm lysate incubated with DTT. Buffer control with DTT and worm lysate without DTT did not show H_2_S production. Data from three independent experiments is shown. The H_2_S levels reference chart is shown on the right side. (B) Model for the mechanism of activation of the hypoxia response pathway by DTT.

## Discussion

Studies in the early 1990s indicated that DTT affects the oxidative environment in the ER without impacting other cellular processes (Lodish & Kong, 1993; Jamsa *et al*, 1994; Braakman *et al*, 1992; Tatu *et al*, 1993). Since then, DTT has been used as an ER-specific stressor in a large number of studies in various model organisms (Pollard *et al*, 1998; Oslowski & Urano, 2011; Qin *et al*, 2010; Frand & Kaiser, 1998; Merksamer *et al*, 2008). Meanwhile, some studies suggested that DTT might broadly impact cellular physiology and affect cellular processes beyond ER (Sims *et al*, 2005; Guillemette *et al*, 2007; MacKenzie *et al*, 2005; Gokul & Singh, 2022; Winter *et al*, 2022). However, the mechanisms of such physiological impacts of DTT remained unexplored. Our current study shows that DTT activates the hypoxia response pathway and challenges the current model that DTT disrupts only the ER milieu in the cell. Our study also highlights that some of the earlier work that used DTT as an ER-specific stressor might need reevaluation in light of the role of DTT in activating the hypoxia response pathway.

Thiol antioxidants result in the activation of futile oxidative cycles in the ER, enhancing disulfide bond turnover (Maity *et al*, 2016). Disulfide bond formation consumes oxygen and produces hydrogen peroxide (Zito, 2015). However, neither oxygen consumption nor hydrogen peroxide production seems to be required for DTT-mediated hypoxia response pathway activation. Earlier studies have shown that oxidative stress could activate the HIF-1-dependent hypoxia response (Lee *et al*, 2010). However, we did not observe the upregulation of *rips-1* under oxidative stress conditions. Because different stimulants could result in non-overlapping responses via HIF-1 (Miller *et al*, 2011), it is possible that the oxidative stress-triggered hypoxia response does not involve the upregulation of *rips-1*. However, the DTT-mediated hypoxia response pathway involves the upregulation of *rips-1*, suggesting that DTT-induced activation of the hypoxia response pathway is unlikely because of oxidative stress.

We discovered that DTT results in H_2_S production when incubated with worm lysate. H_2_S is known to activate the hypoxia response pathway via CYSL-1 (Ma *et al*, 2012). Because DTT and H_2_S activate the hypoxia response through the same pathway, DTT may activate the hypoxia pathway via H_2_S production (Fig 6B). However, from our current data, it is not possible to determine whether the amount of H_2_S produced by DTT is sufficient to induce the hypoxia response pathway. It is also possible that DTT directly acts on CYSL-1, similar to H_2_S (Ma *et al*, 2012), to activate the hypoxia response pathway. Our studies do not rule out the possibility of other mechanisms of activation of the hypoxia response pathway by DTT. It is important to note that reduced thiols in the cell could also enhance HIF-1 degradation by enhancing the activity of EglN1 (Briggs *et al*, 2016), the *C. elegans egl-9* homolog. Therefore, thiol antioxidants could have opposing roles in stabilizing HIF-1, and the outcome might depend on the context or the amount of reduced thiols. Future studies exploring the precise mechanisms of activation of the hypoxia response pathway by DTT in different model systems, including human cell lines, will be very useful in understanding the link between thiol reductive stress and the hypoxia response pathway.

Despite links with various pathological conditions, the physiological effects of reductive stress remain poorly characterized (Rajasekaran, 2020; Ma *et al*, 2020; Pérez-Torres *et al*, 2017; Handy & Loscalzo, 2017). In this study, we show that thiol reductive stress activates the hypoxia response pathway. The activation of the hypoxia response pathway appears to have opposing roles in thiol toxicity in *C. elegans*. By enhancing the expression of *rips-1* under thiol reductive stress, the hypoxia response pathway predisposes *C. elegans* to thiol-mediated toxicity (Fig 6B). Interestingly, SAM-dependent methyltransferases, that are induced by DTT, confer protection against thiol stress in some organisms (MacKenzie *et al*, 2005; Owens *et al*, 2015; Manzanares-miralles *et al*, 2016; Dolan *et al*, 2014). Therefore, the activation of *rips-1* might be a defense response against thiol stress and its toxic effects because of SAM depletion (Gokul & Singh, 2022) could be a side effect. The hypoxia response pathway is known to activate antioxidant responses (Vora *et al*, 2022), which may enhance thiol toxicity. However, because different stimulants could result in non-overlapping responses via HIF-1 (Miller *et al*, 2011), comparing the HIF-1 targets activated by hypoxia vs. thiol stress will be important. The hypoxia response pathway appears to protect against thiol reductive stress independent of *rips-1* expression (Fig 6B). Hypoxia has been shown to protect against mitochondrial reductive stress by enhancing the production of L-2-hydroxyglutarate (Oldham *et al*, 2015). The hypoxia response pathway also enhances the catabolism of hydrogen sulfide (H_2_S) by enhancing the expression of sulfide:quinone oxidoreductase (Budde & Roth, 2011; Malagrinò *et al*, 2019). In future studies, understanding the players downstream of HIF-1 that protect against thiol stress would help establish the interactions between thiol reductive stress and the hypoxia response pathway.

## Materials and Methods

### Bacterial strains

The following bacterial strains were used in the current study: *Escherichia coli* OP50, *E. coli* HT115(DE3), *Pseudomonas aeruginosa* PAO1, and *Comamonas aquatica* DA1877. The cultures of *E. coli* OP50, *E. coli* HT115(DE3), and *C. aquatica* DA1877 were grown in Luria-Bertani (LB) broth at 37°C. The *P. aeruginosa* PAO1 cultures were grown in brain heart infusion (BHI) broth at 37°C.

### *C. elegans* strains and growth conditions

*C. elegans* hermaphrodites were maintained at 20°C on nematode growth medium (NGM) plates seeded with *E. coli* OP50 as the food source unless otherwise indicated. Bristol N2 was used as the wild-type control unless otherwise indicated. The following strains were used in the study: JSJ13 *jsnIs1[rips-1p::GFP + myo-2p::mCherry]*, ZG31 *hif-1(ia4)*, RE666 *ire-1(v33)*, *xbp-1(tm2482)*, RB772 *atf-6(ok551)*, RB545 *pek-1(ok275)*, VC3201 *atfs-1(gk3094)*, and CL2166 *dvIs19[(pAF15)gst-4p::GFP::NLS]*. The following strains were generated in this study: JSJ14 *jsnIs1[rips-1p::GFP + myo-2p::mCherry];rhy-1(jsn13);rhy-1(jsn14)*, JSJ15 *jsnIs1[rips-1p::GFP + myo-2p::mCherry];egl-9(jsn15)*, JSJ16 *jsnIs1[rips-1p::GFP + myo-2p::mCherry];vhl-1(jsn16)*, JSJ17 *jsnIs1[rips-1p::GFP + myo-2p::mCherry];cysl-1(jsn17)*, and JSJ18 *jsnIs1[rips-1p::GFP + myo-2p::mCherry];cysl-1(jsn18)*. Some of the strains were obtained from the Caenorhabditis Genetics Center (University of Minnesota, Minneapolis, MN). The *hif-1(ia4);rips-1p::GFP*, *ire-1(v33);rips-1p::GFP*, *xbp-1(tm2482);rips-1p::GFP*, *atf-6(ok551);rips-1p::GFP*, *pek-1(ok275);rips-1p::GFP*, and *atfs-1(gk3094);rips-1p::GFP* strains were obtained by standard genetic crosses.

### Preparation of NGM plates with different supplements

The following supplements with their product numbers were obtained from HiMedia BioSciences: DTT (#RM525), vitamin B12 (cyanocobalamin) (#PCT0204), ferric chloride (#TC583), pyridoxine hydrochloride (#TC039), NAC (#RM3142) and tert-butyl hydroperoxide (#RM2022). Paraquat (#856177) and MTT (#M5655) were obtained from Sigma-Aldrich. The supplements for *C. elegans* exposure were added to NGM just before pouring them into plates to obtain the desired final concentration.

### *C. elegans* development assays

Synchronized *C. elegans* eggs were obtained by transferring 20-25 gravid adult hermaphrodites on NGM plates for egg-laying for 2 hours. After 2 hours, the gravid adults were removed. The synchronized eggs were incubated at 20°C for 72 hours. After that, the animals in different development stages (L1/L2, L3, and L4/adult) were quantified. Representative images of the NGM plates at the time of quantification of development were also captured. At least three independent experiments (biological replicates) were performed for each condition.

### RNA interference (RNAi)

RNAi was used to generate loss-of-function RNAi phenotypes by feeding worms with *E. coli* HT115(DE3) expressing double-stranded RNA homologous to a target *C. elegans* gene. *E. coli* with the appropriate vectors were grown in LB broth containing ampicillin (100 μg/mL) at 37°C overnight and plated onto RNAi NGM plates containing 100 μg/mL ampicillin and 3 mM isopropyl β-D-thiogalactoside (IPTG). The RNAi-expressing bacteria were allowed to grow overnight at 37°C on RNAi plates. The worms were synchronized on RNAi plates, and the eggs were allowed to develop at 20°C for 72 hours. The RNAi clones were from the Ahringer RNAi library and were verified by sequencing.

### Forward genetic screens for mutants with upregulated *rips-1p::GFP* expression levels

Ethyl methanesulfonate (EMS) mutagenesis screens (Singh, 2021) were performed using the *rips-1p::GFP* strain. Approximately 2500 synchronized late L4 larvae were treated with 50 mM EMS for 4 hours and then washed three times with M9 medium. The washed animals (P0 generation) were then transferred to 9-cm NGM plates containing *E. coli* OP50 and allowed to lay eggs (F1 progeny) overnight. The P0s were then washed away with M9 medium, while the F1 eggs remained attached to the bacterial lawn. The F1 eggs were allowed to grow to adulthood. The adult F1 animals were bleached to obtain F2 eggs. The F2 eggs were transferred to *E. coli* OP50 plates containing 50 nM vitamin B12 and incubated at 20°C for 72 hours. After that, the plates were screened for animals that had high GFP levels. Approximately 50,000 haploid genomes were screened, and three mutants were isolated. All of the mutants were backcrossed six times with the parental *rips-1p::GFP* strain before analysis.

### Forward genetic screens for mutants that lacked DTT-mediated *rips-1p::GFP* upregulation

All the mutagenesis steps were identical to the screen for mutants with upregulated *rips-1p::GFP* expression levels until the F2 eggs stage. The F2 eggs were transferred to *E. coli* OP50 plates and incubated at 20°C for 72 hours. After that, the adult F2 animals were washed away with M9 and transferred to *E. coli* OP50 plates containing 10 mM DTT. After 10 hours of incubation, the plates were screened for animals that lacked GFP induction. Approximately 50,000 haploid genomes were screened, and two mutants were isolated. Both the mutants were backcrossed six times with the parental *rips-1p::GFP* strain before analysis.

### Whole-genome sequencing (WGS) and data analysis

The genomic DNA was isolated as described earlier (Singh & Aballay, 2017; Gokul & Singh, 2022). Briefly, the mutant animals were grown at 20°C on NGM plates seeded with *E. coli* OP50 until starvation. Four 9-cm *E. coli* OP50 plates were used for each strain to obtain a sufficient number of animals. The animals were rinsed off the plates with M9, washed three times, incubated in M9 with rotation for 2 hours to eliminate food from the intestine, and washed three times again with distilled water, followed by storage at -80°C until genomic DNA extraction. Genomic DNA extraction was performed using the Gentra Puregene Kit (Qiagen, Netherlands). DNA libraries were prepared according to a standard Illumina (San Diego, CA) protocol. The DNA was subjected to WGS on an Illumina NovaSeq 6000 sequencing platform using 150 paired-end nucleotide reads. Library preparation and WGS were performed at the National Genomics Core, National Institute of Biomedical Genomics, Kalyani, India.

The whole-genome sequence data were analyzed using the web platform Galaxy as described earlier (Gokul & Singh, 2022). Briefly, the forward and reverse FASTQ reads, *C. elegans* reference genome Fasta file (ce11M.fa), and the gene annotation file (SnpEff4.3 WBcel235.86) were input into the Galaxy workflow. The low-quality ends of the FASTQ reads were trimmed using the Sickle tool. The trimmed FASTQ reads were mapped to the reference genome Fasta files with the BWA-MEM tool. Using the MarkDuplicates tool, any duplicate reads (mapped to multiple sites) were filtered. Subsequently, the variants were detected using the FreeBayes tool that finds small polymorphisms, including single-nucleotide polymorphisms (SNPs), insertions and deletions (indels), multi-nucleotide polymorphisms (MNPs), and complex events (composite insertion and substitution events) smaller than the length of a short-read sequencing alignment. The common variants among different mutants were subtracted. The SnpEff4.3 WBcel235.86 gene annotation file was used to annotate and predict the effects of genetic variants (such as amino acid changes). Finally, the linkage maps for each mutant were generated using the obtained variations.

### *C. elegans* RNA isolation and quantitative reverse transcription-PCR (qRT-PCR)

Animals were synchronized by egg laying. Approximately 40 gravid adult animals were transferred to 9-cm *E. coli* OP50 plates without DTT and allowed to lay eggs for 4 hours. The gravid adults were then removed, and the eggs were allowed to develop at 20°C for 72 hours. Subsequently, the synchronized adult animals were collected with M9, washed twice with M9, and then transferred to 9-cm *E. coli* OP50 plates containing 10 mM DTT. The control animals were maintained on *E. coli* OP50 plates without DTT. After the transfer of the animals, the plates were incubated at 20°C for 10 hours. Subsequently, the animals were collected, washed with M9 buffer, and frozen in TRIzol reagent (Life Technologies, Carlsbad, CA). Total RNA was extracted using the RNeasy Plus Universal Kit (Qiagen, Netherlands). A total of 1 μg of total RNA was reverse-transcribed with random primers using the PrimeScript™ 1st strand cDNA Synthesis Kit (TaKaRa) according to the manufacturer’s protocols. qRT-PCR was conducted using TB Green fluorescence (TaKaRa) on a LightCycler 480 II System (Roche Diagnostics) in 96-well-plate format. Twenty microliter reactions were analyzed as outlined by the manufacturer (TaKaRa). The relative fold-changes of the transcripts were calculated using the comparative *CT*(2^-ΔΔ*CT*^) method and normalized to pan-actin (*act-1, -3, -4*) as described earlier (Singh & Aballay, 2019a). All samples were run in triplicate and repeated at least three times. The sequence of the primers is provided in Table S2.

### Fast-killing assays on *P. aeruginosa* PAO1

For fast-killing assays, *P. aeruginosa* PAO1 cultures were grown by inoculating individual bacterial colonies into 4-5 mL of BHI broth and incubated for 10–12 hours on a shaker at 37°C. Then, the cultures were diluted ten times in BHI broth, and 50 μL of the diluted culture was spread on a 3.5-cm BHI agar plate. After spreading bacteria, plates were incubated at 37°C for 24 hours. After that, the plates were allowed to cool to room temperature, and worms synchronized on NGM plates seeded with *C. aquatica* DA1877 with and without 5 mM DTT at the L4 stage were transferred to them. Plates were kept at room temperature and scored for live and dead worms at indicated times of transfer.

### H_2_S production assay

The H_2_S production capacity assay was adapted from an earlier study (Statzer *et al*, 2022) and modified for measuring H_2_S production by DTT. Briefly, synchronized wild-type N2 young adult *C. elegans* hermaphrodites were washed off the plates with M9. The worms were washed three times with M9 to remove residual bacteria. The semisoft pellet of worms was weighed to ensure an equal number of worms in each sample (100 µg of worm pellet per sample) and mixed with 800 µL of 1× passive lysis buffer (Promega, #E1910). This was followed by eight cycles of sonication with 3 s on and 30 s off at an amplitude of 40%. The suspension was centrifuged at 12,000 × *g* at 4°C for 10 min. The supernatant (800 µL) was transferred to new microcentrifuge tubes. The fuel mix was prepared by adding 20 mM DTT and 2.5 mM pyridoxine hydrochloride in phosphate buffer saline. The fuel mix (200 µL) was added to the worm lysate to make the final DTT and pyridoxine hydrochloride concentrations 4 mM and 0.5 mM, respectively. The tubes containing worm lysate and DTT were sealed with H_2_S sensing lead acetate strips (HiMedia BioSciences # WT041) and incubated at 37°C for 12 hours. After 12 hours of incubation, lead acetate strips were observed to determine H_2_S production.

### Impact of oxidative stress on *rips-1p::GFP* expression

Egg laying of *rips-1p::GFP* and *gst-4p::GFP* animals was carried out by transferring gravid adult hermaphrodites on NGM plates seeded with *E. coli* OP50 for 4 hours. Worms were removed after egg laying, and the plates containing eggs were incubated at 20°C. After 72 hours, synchronized adult worms were transferred to *E. coli* OP50 plates containing 1 mM paraquat, 5 mM tert-butyl hydroperoxide, 10 mM DTT, and 7.5 mM ferric chloride for 12 hours. After that, the animals were prepared for fluorescence imaging.

### Fluorescence imaging of *C. elegans*

Fluorescence imaging was carried out as described previously (Gokul & Singh, 2022; Singh & Aballay, 2019b). Briefly, the fluorescence reporter strains were picked using a non-fluorescence stereomicroscope to avoid potential bias. The animals were anesthetized using an M9 salt solution containing 50 mM sodium azide and mounted onto 2% agarose pads. The animals were then visualized using a Nikon SMZ-1000 fluorescence stereomicroscope. The fluorescence intensity was quantified using Image J software.

### Cell culture

Human cell lines, HeLa (cervical cancer), LN229 (glioblastoma), and Cal27 (oral squamous cell carcinoma), were obtained from the National Center of Cell Science (NCCS), Pune. The cells were cultured in a humidified incubator at 37°C and 5% CO_2_ in Dulbecco’s modified eagle medium (DMEM), supplemented with 10% fetal bovine serum (FBS). The culture medium was replenished every third day. The study with human cell lines was approved by the institute ethics committee, PGIMER, with approval number IEC-01/2023-2648.

### MTT cell viability assay

To assess cell viability, HeLa, LN229, and Cal27 cell lines were exposed to 10 mM DTT for 24 hours, followed by measurement of their survival using MTT (3-(4,5-Dimethylthiazol-2-yl)-2,5-Diphenyltetrazolium Bromide) assay as reported previously (Kumar *et al*, 2017, 2021). Briefly, the cells were seeded at an initial density of 5×10^3^ per well in 96-well cell culture plates and treated with 10 mM DTT for 24 hours while the control cells remained untreated. After that, the cells were washed to remove DTT. Then, the cells were treated with 10% MTT solution for four hours in the dark. The formazan crystals formed from MTT by live cells were solubilized by DMSO. The absorbance of the solution was measured at 570 nm wavelength on a microplate reader (Infinite 200 Pro, TECAN). Readings from 12 different wells were obtained for each condition.

### Cell lines RNA isolation and qRT-PCR

HeLa, LN229, and Cal27 cell lines were treated with 10 mM DTT for 24 hours. The total RNA of DTT treated and control cells was isolated using the TRIzol method. Total RNA was reverse transcribed to cDNA using the High-Capacity cDNA Reverse Transcription Kit (Applied Biosystems), as per the manufacturer’s instructions. qRT-PCR was conducted using DyNAmo ColorFlash SYBR Green (ThermoFisher Scientific) on a CFX96 Real-Time System (Bio-Rad) in 96-well-plate format. Twenty microliter reactions were analyzed as outlined by the manufacturer (Bio-Rad). The relative fold changes of the transcripts were calculated using the comparative *CT*(2^-ΔΔ*CT*^) method and normalized to beta-actin. All samples were run in triplicate and repeated at least three times. The sequence of the primers is provided in Table S2.

### Hypoxia staining of human cell lines and quantification

Human cell lines were pretreated with 10 mM DTT for 24 hours. After that, DTT was removed, and the cells were treated with Image-iT™ Green Hypoxia Reagent (Invitrogen) at a concentration of 1 µM for 30 minutes at 37°C. The reagent was then removed, and the cells were incubated in either DMEM (control) or DMEM containing 10 mm DTT for 4 hours. After that, cell images were captured using a fluorescence microscope (Leica) at the excitation/emission wavelength of 488/520 nm. Simultaneously, cells were separately cultured in 96-well plates and stained with the Image-iT™ Green Hypoxia Reagent as described above. The fluorescence of the cells was then measured in a fluorescence plate reader (Infinite 200 Pro, TECAN) at the excitation/emission wavelengths of FITC (488/520 nm) with the following parameters: Z-position 20000 µm, gain 100, excitation bandwidth 9 nm, emission bandwidth 20 nm, and the number of flashes 25. As a control for the redox environment, the cells were treated with 10 mM NAC for 24 hours and stained with the Image-iT™ Green Hypoxia Reagent, like the treatment with DTT. The pH of NAC was adjusted to 7.0 using a sodium hydroxide solution (NaOH). The fluorescence of the cells was measured in a fluorescence plate reader as described for DTT.

### Oxygen consumption rate (OCR) measurement

Human cell lines HeLa, LN229, and Cal27 were seeded at a density of 2×10^4^ cells per well in a Seahorse XFe24 Cell Culture Microplate (102340-100; Agilent, Santa Clara, CA). Cells were incubated overnight for adherence in a culture media composed of DMEM and 10% FBS at 37°C and 5% CO_2_. Before OCR measurements, the cells were washed twice in DMEM, devoid of sodium bicarbonate (pH adjusted with NaOH), and incubated in a BOD incubator at 37°C for 1 hour. The cells were in DMEM without FBS and sodium bicarbonate at a final volume of 450 μl per well. The culture medium controls contained only the medium without cells. The plate with cells was then placed into a Seahorse XFe24 Analyzer (Agilent, Santa Clara, CA), and the OCR measurements were carried out at 37°C. The sensor cartridge was hydrated overnight before using for OCR measurements. After three readings for basal OCR, 50 µL of 100 mM DTT was injected into different wells to obtain a final concentration of 10 mM DTT. For 0 mM DTT controls, 50 µL of water was injected into wells. Three technical replicates were set for each condition in each experiment. Two independent experiments were carried out.

### Quantification and Statistical Analysis

The statistical analysis was performed with Prism 8 (GraphPad). All error bars represent mean±standard deviation (SD) unless otherwise indicated. The two-sample *t*-test was used when needed, and the data were judged to be statistically significant when p < 0.05. In the figures, asterisks (*) denote statistical significance as follows: *, p < 0.05, **, p < 0.01, ***, p < 0.001, as compared with the appropriate controls.

## Data Availability

The whole-genome sequence data for JSJ14-JSJ18 have been submitted to the public repository, the Sequence Read Archive, with BioProject ID PRJNA933286.

## Acknowledgments

We thank Shweta Saini (IISER Mohali) for assistance with *C. elegans* forward genetic screens and Aditi Bhagat (PGIMER Chandigarh) for assistance with cell culture experiments. Some strains used in this study were provided by the Caenorhabditis Genetics Center (CGC), which is funded by the NIH Office of Research Infrastructure Programs (P40 OD010440). This work was supported by the Ramalingaswami Re-entry Fellowship (Ref. No. BT/RLF/Re-entry/50/2020) awarded by the Department of Biotechnology, India, Science and Engineering Research Board (SERB) Startup Research Grant (Ref. No. SRG/2020/000022) awarded by DST, India, and IISER Mohali intramural funds.

## Author contributions

Ravi and J.S. conceived and designed the study; Ravi performed the *C. elegans* experiments; Ravi and J.S. analyzed and interpreted the *C. elegans* data; A.K. planned, performed, and analyzed human cell line experiments; S.B. supervised human cell line experiments; J.S. wrote the manuscript with inputs from Ravi and A.K.

## Expanded View Figure Legends

**Figure S1.**
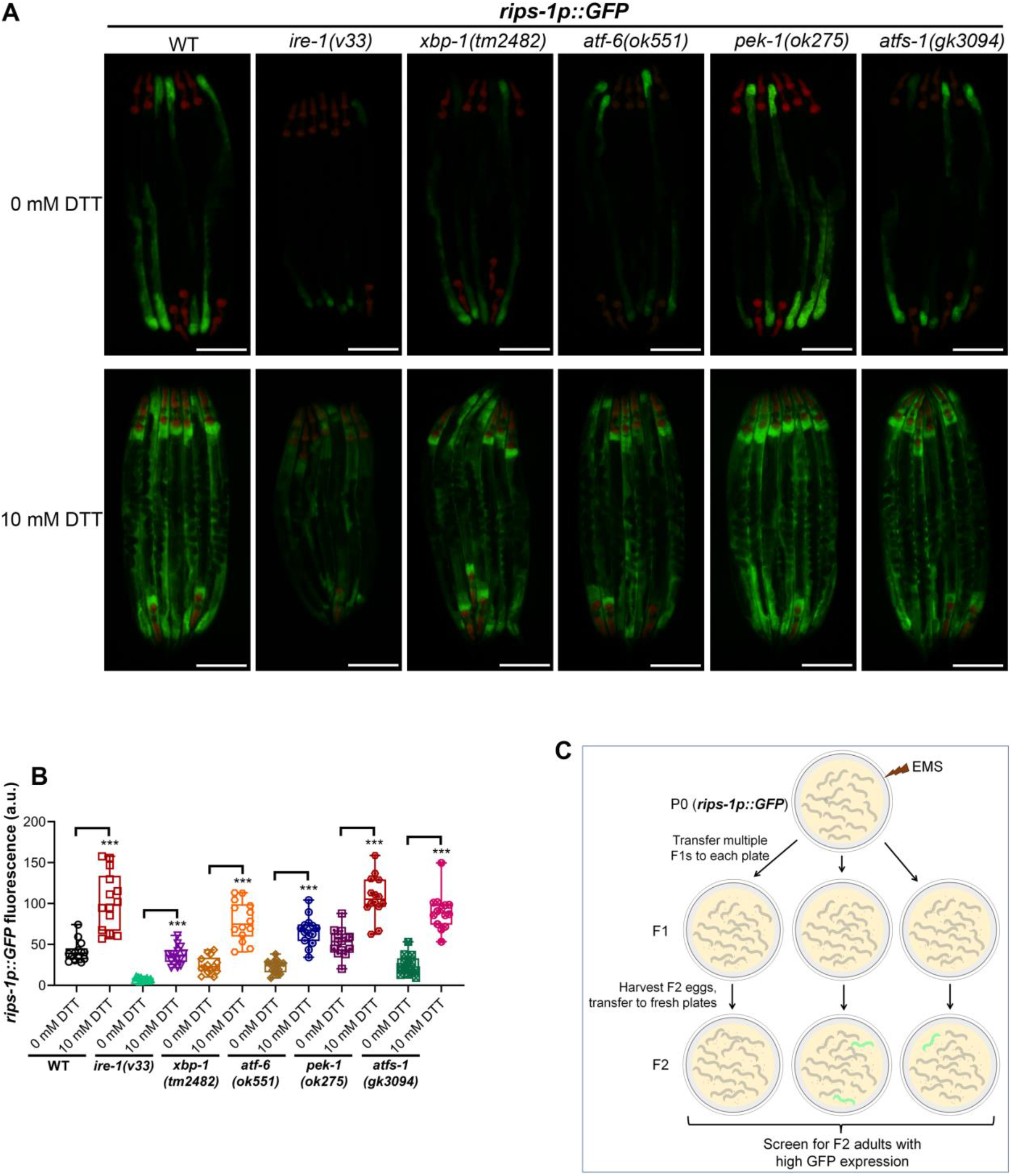
The unfolded protein response (UPR) pathways in the endoplasmic reticulum (ER) and mitochondria do not control DTT-mediated *rips-1* upregulation. (A) Representative fluorescence images of *rips-1p::GFP* in the background of WT, *ire-1(v33)*, *xbp-1(tm2482)*, *atf-6(ok551)*, *pek-1(ok275)*, and *atfs-1(gk3094)* animals grown on 0 mM DTT until the young adult stage, followed by incubation on 0 mM or 10 mM DTT for 10 hours. The red fluorescence in the pharynx region is from the *myo-2p::mCherry* coinjection marker. Scale bar = 200 µm. (B) Quantification of GFP levels of *rips-1p::GFP* in the background of WT, *ire-1(v33)*, *xbp-1(tm2482)*, *atf-6(ok551)*, *pek-1(ok275)*, and *atfs-1(gk3094)* animals grown on 0 mM DTT until the young adult stage, followed by incubation on 0 mM or 10 mM DTT for 10 hours. ***p < 0.001 via the *t*-test (*n* = 14 worms each). (C) Scheme for a forward genetic screen for mutants with constitutive upregulation of *rips-1p::GFP*.

**Figure S2.**
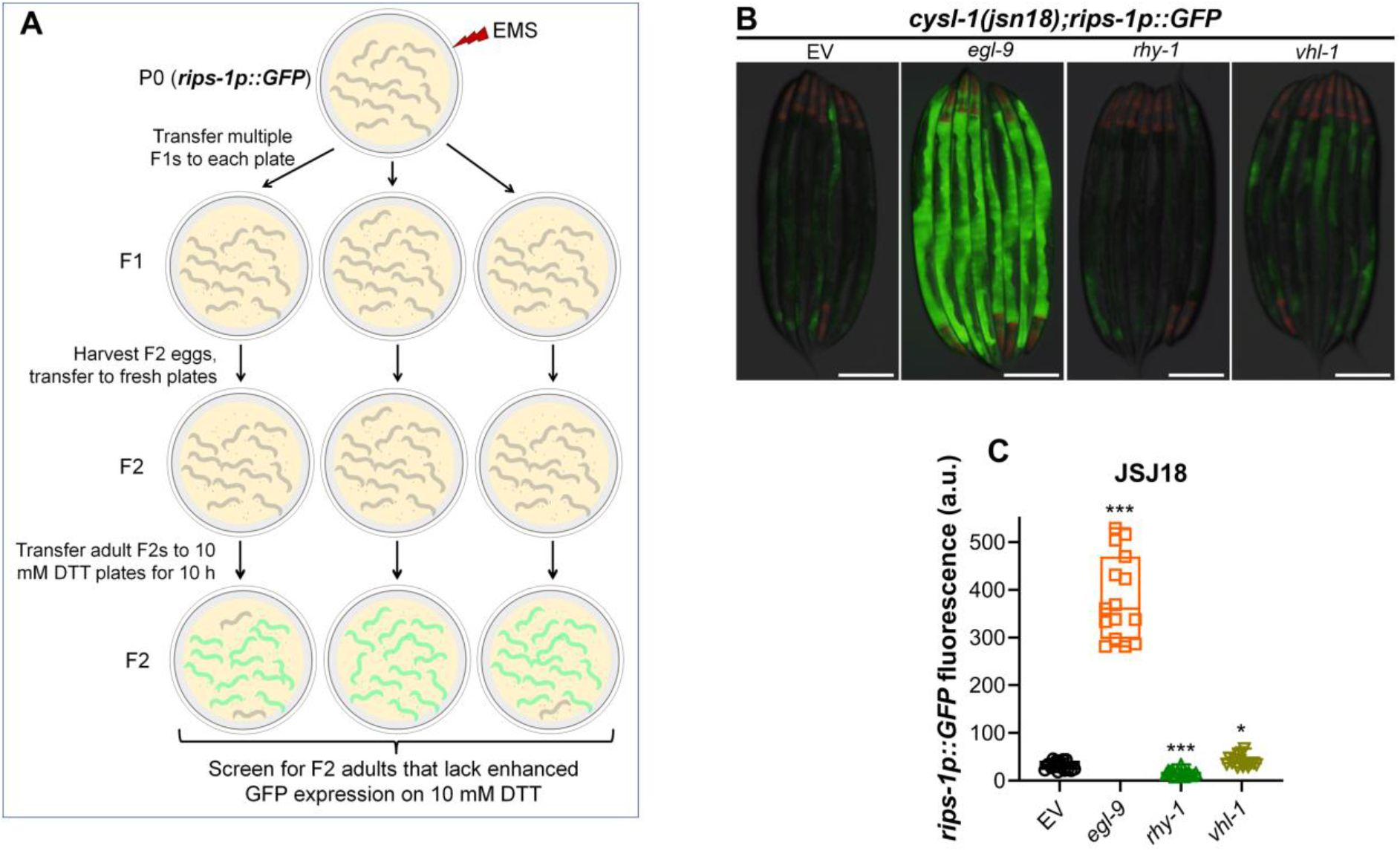
CYSL-1 is required for DTT-mediated *rips-1* upregulation. (A) Scheme for a forward genetic screen for mutants that lack upregulation of *rips-1p::GFP* on 10 mM DTT. (B) Representative fluorescence images of JSJ18 (*cysl-1(jsn18);rips-1p::GFP*) animals following RNAi against *egl-9*, *rhy-1*, and *vhl-1*, along with EV control. Scale bar = 200 µm. (C) Quantification of GFP levels of JSJ18 (*cysl-1(jsn18);rips-1p::GFP*) animals following RNAi against *egl-9*, *rhy-1*, and *vhl-1*, along with EV control. ***p < 0.001 and *p < 0.05 via the *t*-test (*n* = 15 worms each).

**Figure S3.**
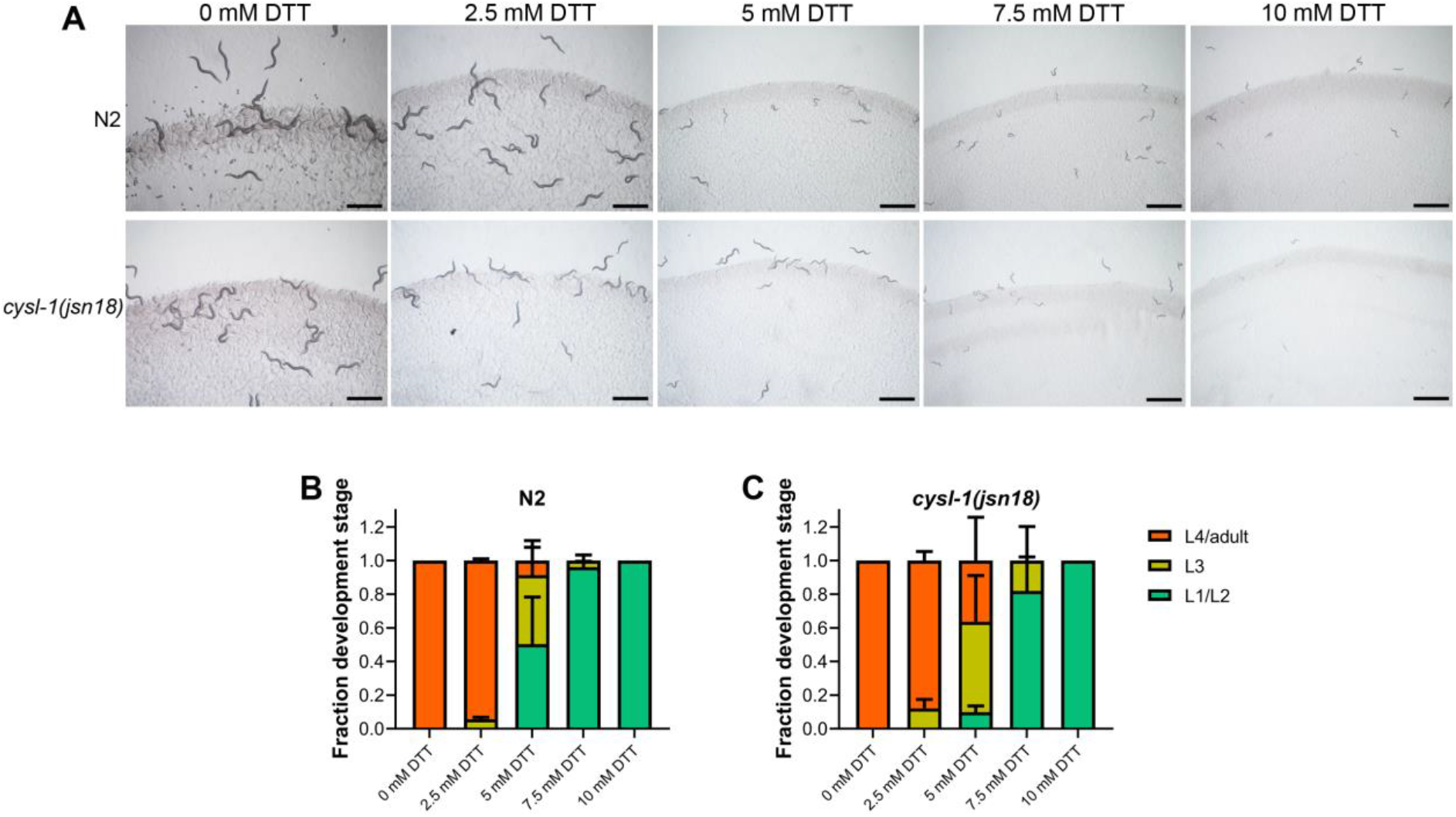
*cysl-1(jsn18)* animals develop better than N2 animals at 5 mM DTT. (A) Representative images of N2 and *cysl-1(jsn18)* animals on various concentrations of DTT on *E. coli* OP50 diet after 72 hours of hatching at 20°C. Scale bar = 1 mm. (B-C) Quantification of different developmental stages of N2 (B) and *cysl-1(jsn18)* (C) animals on various concentrations of DTT on *E. coli* OP50 diet after 72 hours of hatching at 20°C. The N2 data in (B) are the same as in Fig 4B. (*n* = 3 biological replicates; animals per condition per replicate >100).

**Figure S4.**
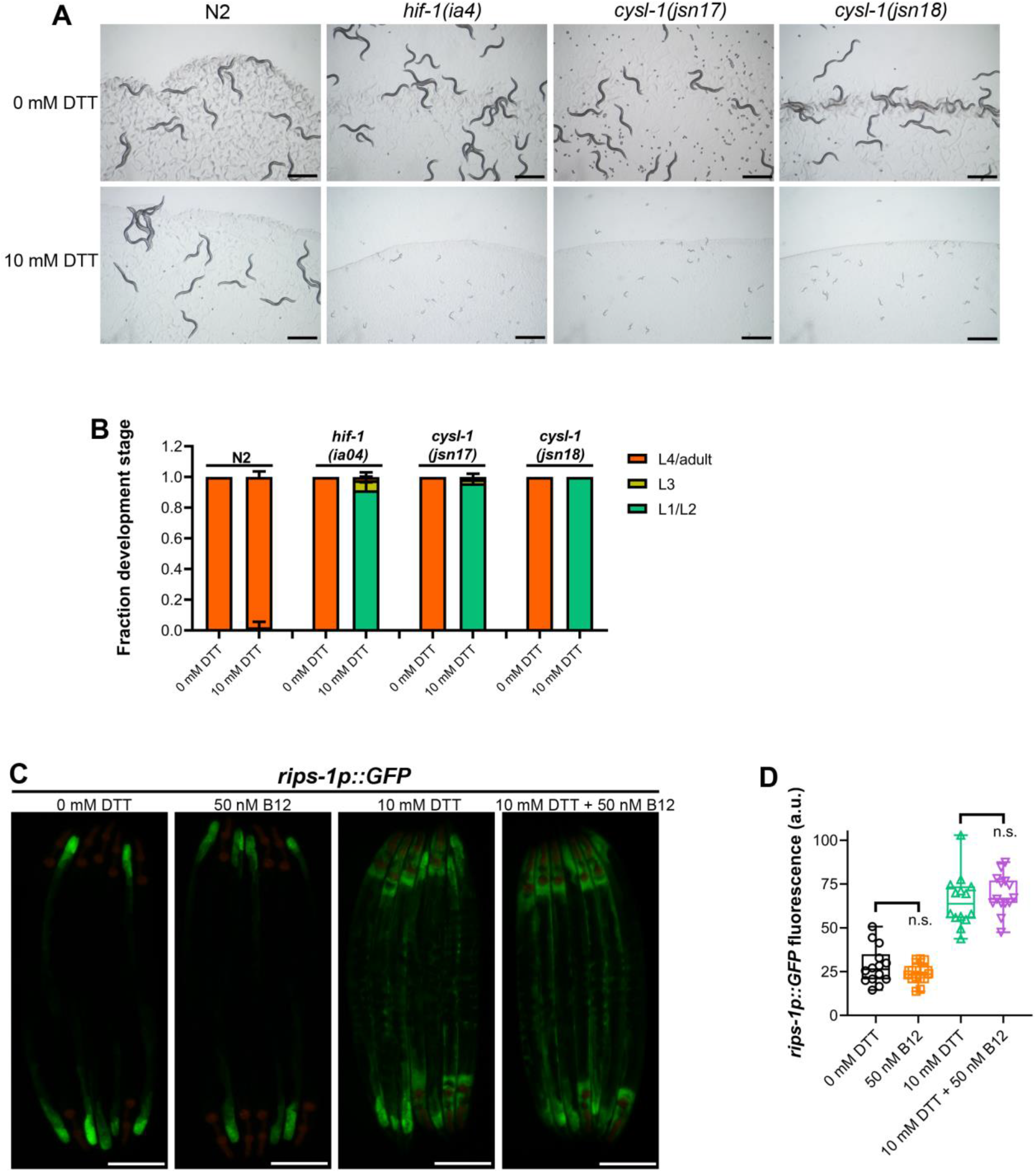
Hypoxia response pathway protects against thiol reductive stress. (A) Representative images of N2, *hif-1(ia4)*, *cysl-1(jsn17)*, and *cysl-1(jsn18)* animals after 72 hours of hatching at 20°C on *C. aquatica* DA1877 diet containing 0 mM or 10 mM DTT. Scale bar = 1 mm. (B) Quantification of different developmental stages of N2, *hif-1(ia4)*, *cysl-1(jsn17)*, and *cysl-1(jsn18)* animals after 72 hours of hatching at 20°C on *C. aquatica* DA1877 diet containing 0 mM or 10 mM DTT. (*n* = 3 biological replicates; animals per condition per replicate >100). (C) Representative fluorescence images of *rips-1p::GFP* animals grown on *E. coli* OP50 diet without any supplements until the young adult stage, followed by incubation on *E. coli* OP50 diet containing no supplement (0 mM DTT), 50 nM vitamin B12, 10 mM DTT, and 10 mM DTT + 50 nM vitamin B12 for 10 hours. Scale bar = 200 µm. (D) Quantification of GFP levels of *rips-1p::GFP* animals grown on *E. coli* OP50 diet without any supplements until the young adult stage, followed by incubation on *E. coli* OP50 diet containing no supplement (0 mM DTT), 50 nM vitamin B12, 10 mM DTT, and 10 mM DTT + 50 nM vitamin B12 for 10 hours. n.s., nonsignificant via the *t*-test (*n* = 14 animals each).

**Figure S5.**
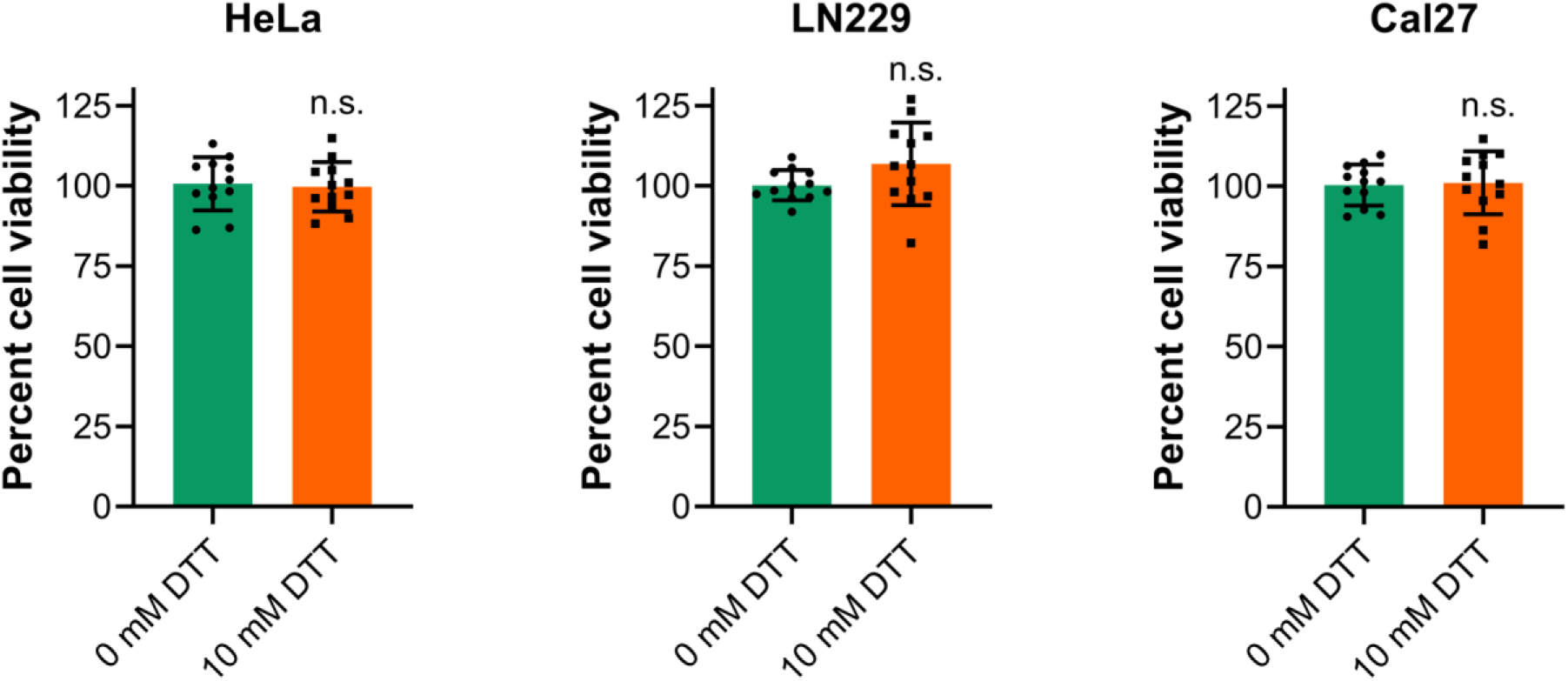
The cell viability is not altered at 10 mM dithiothreitol (DTT). Measurement of percent cell viability using MTT assays in HeLa, LN229, and Cal27 cells upon exposure to 10 mM DTT for 24 hours, along with controls without DTT. n.s., nonsignificant via the *t*-test (*n* = 12 wells each).

**Figure S6.**
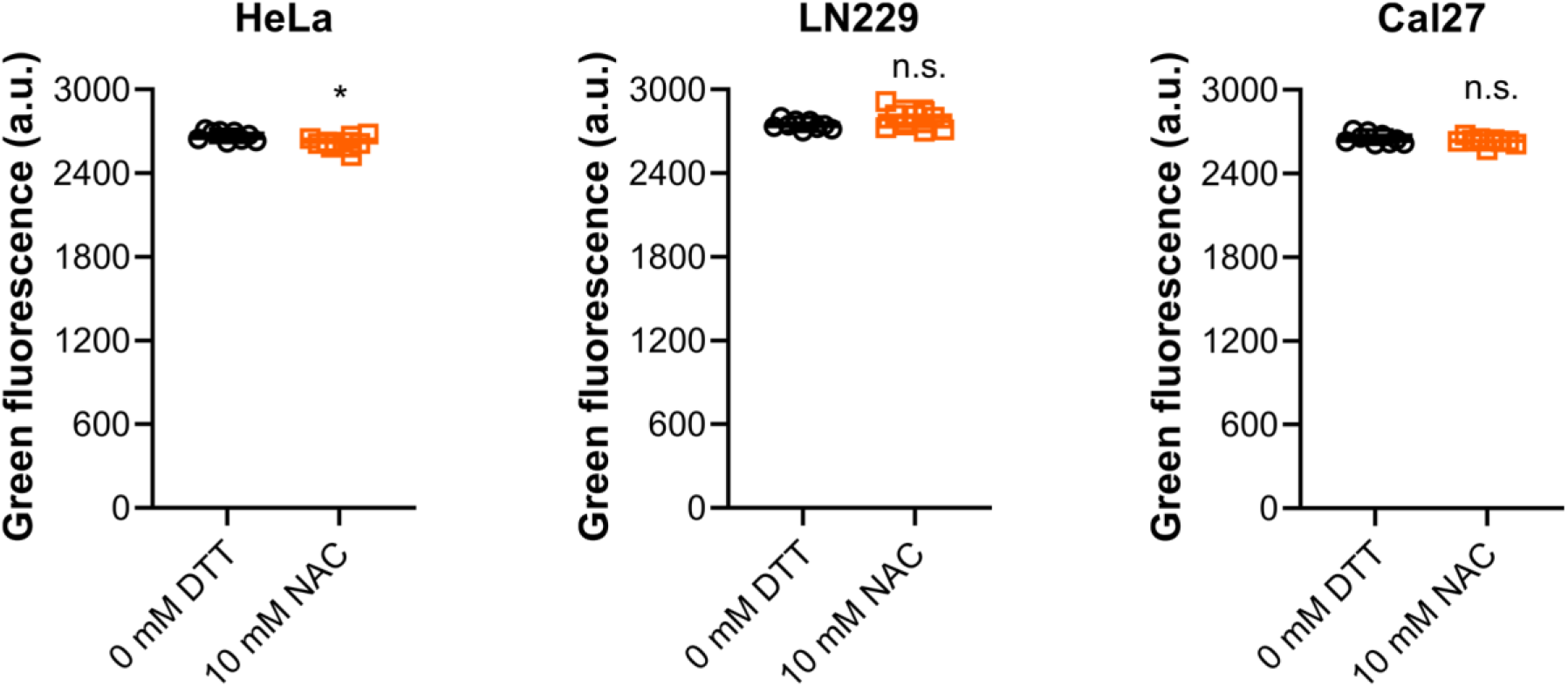
*N*-acetylcysteine (NAC) treatment does not enhance Image-iT™ green hypoxia reagent fluorescence in human cell lines. Quantification of fluorescence levels of Image-iT™ green hypoxia reagent-treated HeLa, LN229, and Cal27 cell lines. The cells were pretreated with 0 mM or 10 mM NAC for 24 hours before treatment with image-iT™ green hypoxia reagent. *p < 0.05 via the *t*-test. n.s., nonsignificant (*n* = 9 wells each).

**Figure S7.**
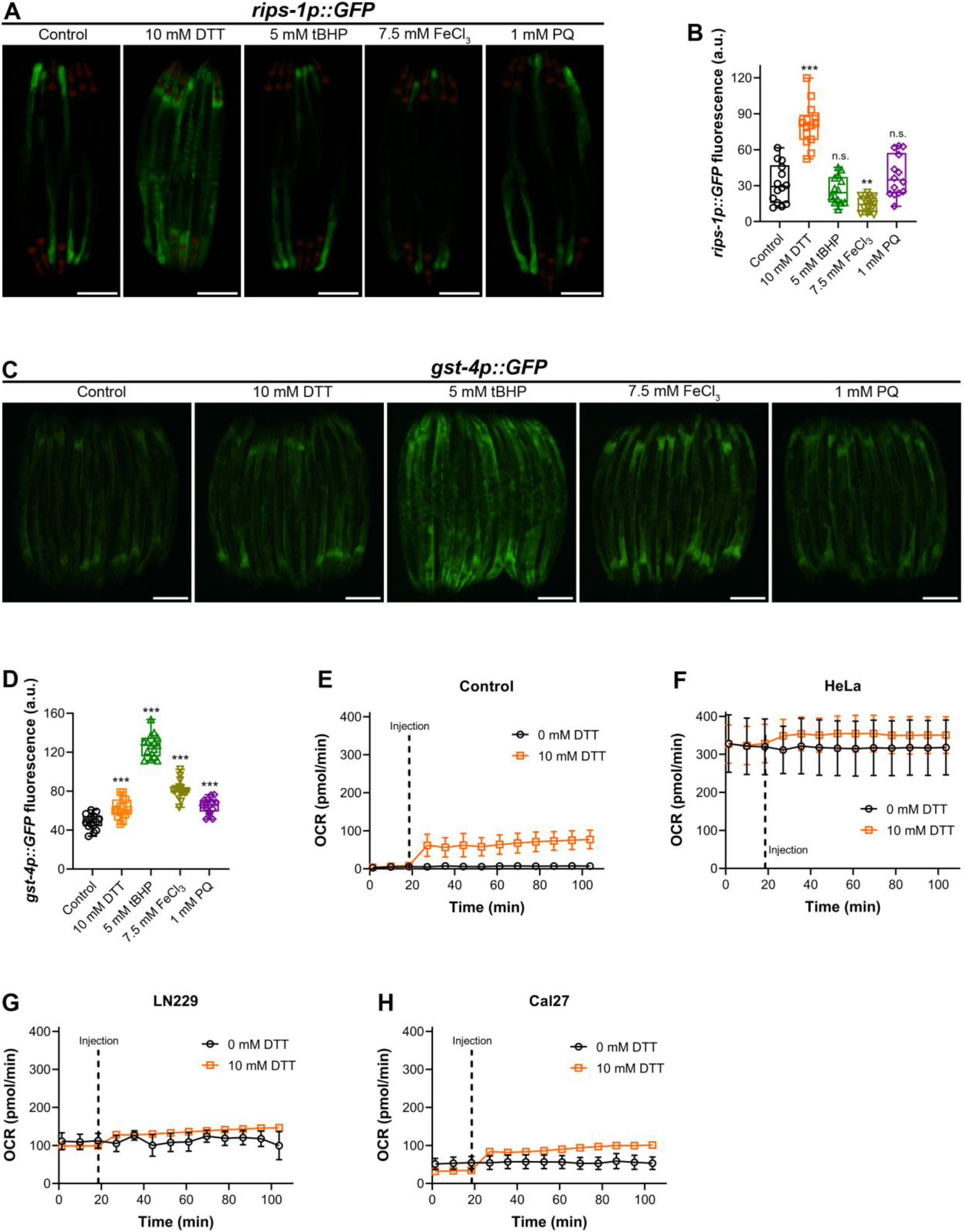
The futile oxidative cycle in the ER under DTT stress does not result in the activation of the hypoxia response pathway. (A) Representative fluorescence images of *rips-1p::GFP* animals grown on *E. coli* OP50 diet without any supplements until the young adult stage, followed by incubation on *E. coli* OP50 diet containing no supplement (control), 10 mM DTT, 5 mM tert-butyl hydroperoxide (tBHP), 7.5 mM ferric chloride (FeCl_3_), and 1 mM paraquat (PQ) for 12 hours. Scale bar = 200 µm. (B) Quantification of GFP levels of *rips-1p::GFP* animals grown on *E. coli* OP50 diet without any supplements until the young adult stage, followed by incubation on *E. coli* OP50 diet containing no supplement (control), 10 mM DTT, 5 mM tBHP, 7.5 mM FeCl_3_, and 1 mM paraquat for 12 hours. ***p < 0.001 and **p < 0.01 via the *t*-test. n.s., nonsignificant (*n* = 14 worms each). (C) Representative fluorescence images of *gst-4p::GFP* animals grown on *E. coli* OP50 diet without any supplements until the young adult stage, followed by incubation on *E. coli* OP50 diet containing no supplement (control), 10 mM DTT, 5 mM tBHP, 7.5 mM FeCl_3_, and 1 mM paraquat (PQ) for 12 hours. Scale bar = 200 µm. (D) Quantification of GFP levels of *gst-4p::GFP* animals grown on *E. coli* OP50 diet without any supplements until the young adult stage, followed by incubation on *E. coli* OP50 diet containing no supplement (control), 10 mM DTT, 5 mM tBHP, 7.5 mM FeCl_3_, and 1 mM paraquat for 12 hours. ***p < 0.001 via the *t*-test (*n* = 15 worms each). (E-H) Oxygen consumption rates (OCR) in medium control without cells (E), HeLa (F), LN229 (G), and Cal27 (H) cells. After three OCR measurements, water or DTT was injected in the wells to obtain 0 mM or 10 mM DTT and is indicated as injection in the plots. The cells were seeded at 2 ×10^4^ cells per well. The data represent mean and standard error from two-three wells per condition.

**Table S1:** Summary of the alleles identified by whole-genome sequencing in JSJ14-JSJ18 strains.

| Strain name | Allele | Mutation | Change in DNA | Chromosomal position |
| --- | --- | --- | --- | --- |
| JSJ14 | *rhy-1(jsn13 jsn14)* | S146Y<br>R183C | tCt/tAt<br>Cgt/Tgt | II, 9162060<br>II, 9162170 |
| JSJ15 | *egl-9(jsn15)* | Splice acceptor variant | G to A | V, 10473679 |
| JSJ16 | *vhl-1(jsn16)* | Q154* | Caa/Taa | X, 11308096 |
| JSJ17 | *cysl-1(jsn17)* | R50C | Cgt/Tgt | X, 9954105 |
| JSJ18 | *cysl-1(jsn18)* | G190E | gGa/gAa | X, 9953478 |

**Table S2:**
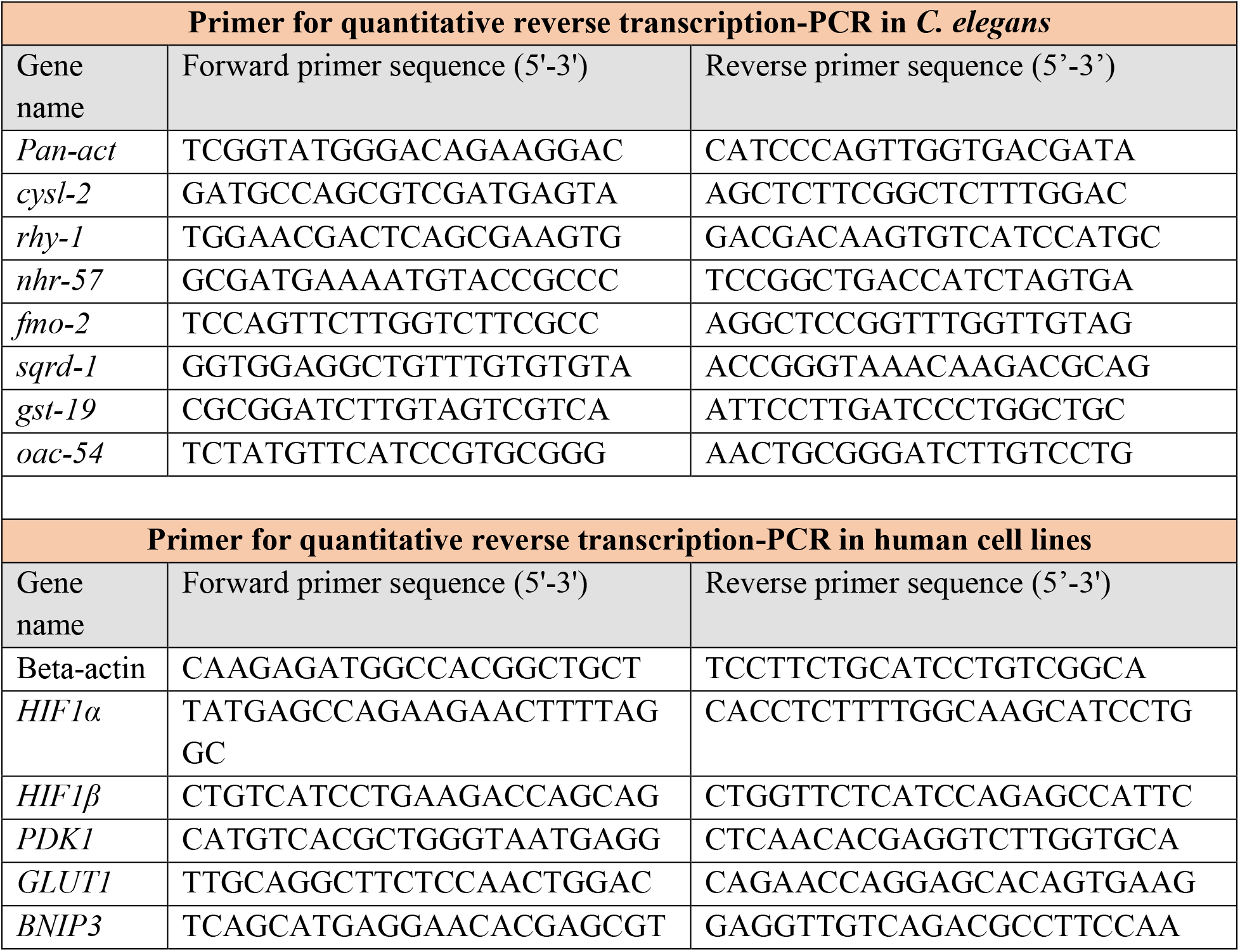
Primers used in the study.

